# TMEM165 acts as a proton-activated Ca^2+^ importer in lysosomes

**DOI:** 10.1101/2023.08.08.552438

**Authors:** Matthew Zajac, Sourajit Mukherjee, Palapuravan Anees, Daphne Oettinger, Katharine Henn, Jainaha Srikumar, Junyi Zou, Anand Saminathan, Yamuna Krishnan

## Abstract

Lysosomal calcium (Ca^2+^) release is critical to cell signaling and is mediated by well-known lysosomal Ca^2+^ channels. Yet, how lysosomes refill their Ca^2+^ remains hitherto undescribed. Here, from an RNAi screen, we identify an evolutionarily conserved gene, *lci-1*, that facilitates lysosomal Ca^2+^ entry in *C. elegans* and in mammalian cells. Its human homolog TMEM165, previously designated as a Golgi-resident Ca^2+^/H^+^ exchanger (CAX), has a minor lysosomal population of unknown function. Using genetics, lysosomal Ca^2+^ imaging and electrophysiology, we show that TMEM165 acts as a proton-activated, lysosomal <u>C</u>a^2+^ importer in lysosomes. Defects in lysosomal Ca^2+^ channels cause several neurodegenerative diseases, and knowledge of lysosomal Ca^2+^ importers may provide new avenues to explore the physiology of Ca^2+^ channels.

## Introduction

Eukaryotic cells use several mechanisms to regulate the spatial and temporal dynamics of cytosolic Ca^2+^[1]. These include feedback with intracellular Ca^2+^ stores such as the endoplasmic reticulum (ER), lysosomes or mitochondria, which have high Ca^2+^ content^1–3^. These organelles harbor proteins that mediate Ca^2+^ release as well as import, because after the organelle releases Ca^2+^ via an exporter, its lumenal Ca^2+^ must be replenished by a Ca^2+^ importer. Thus, after its release via the Ryanodine receptor in the ER, Ca^2+^ is refilled by SERCA^4, 5^ while in the mitochondria, after release via the Na^+^/Ca^2+^ exchanger, NCX, Ca^2+^ is replenished by the mitochondrial Ca^2+^ uniporter, MCU^6–9^. Lysosomes have recently come into prominence as the acidic Ca^2+^ stores of the cell^10^ and while we know of many channels that release Ca^2+^ from lysosomes, no Ca^2+^ importers are known in humans^11^.

Vacuoles perform the functions of lysosomes in plants and are known to harbor two kinds of Ca^2+^ importers: high-capacity Ca^2+^ exchangers and low-capacity Ca^2+^ ATPases^12, 13^. In mammalian lysosomes, both, a pH-dependent Ca^2+^ uptake mechanism and a Ca^2+^/H^+^ exchange mechanism, have been posited^14–16^. The heterologous overexpression of Xenopus CAX in mammalian lysosomes leads to Ca^2+^ import^17^, and a P-type Ca^2+^ ATPase, ATP13A2, facilitates Ca^2+^ accumulation^18^. However, a molecule that directly transports Ca^2+^ into human lysosomes is yet to be found.

## Results

### Targeted screen identifies Y54F10Al.1 as a lysosomal Ca^2+^ importers in nematodes

We performed a targeted RNA interference (RNAi) screen to identify lysosomal Ca^2+^ importers using assays that previously pinpointed ATP13A2 (*catp-6* in *C. elegans*), as facilitating import^17^. Here, gene knockdown reverses the phenotypic defects in a lysosomal Ca^2+^ channel mutant (Figure S1A). In *C. elegans*, deleting *cup-5*, the worm homolog of the lysosomal Ca^2+^ release channel TRPML1, is lethal^19^. We knocked down 228 lysosomal genes in *cup-5*^+/-^ worms (Table S1 not included), screening for reversal of lethality, reasoning that a broken Ca^2+^ intake mechanism would reverse the phenotype. We found that knocking down the uncharacterized gene *Y54F10AL.1*, now denoted *lci-1*, in *cup-5* deficient nematodes rescues lethality (Figure 1A and S1B-D). Next, knocking down *cup-5* leads to abnormally large lysosomes due to storage arising from lysosome dysfunction. We therefore tested whether *lci-1* knockdown rescues the lysosomal *cup-5* phenotype. We used the *arIs37;cup-5(ar465)* strain, because in this *cup-5* hypomorph, lysosomal dysfunction is severe enough to produce swollen, GFP-labeled lysosomes, yet is insufficient for lethality^20^. Indeed, knocking down *lci-1* restores lysosome sizes to normalcy, indicating that lysosome function is restored, potentially because aberrant lysosomal Ca^2+^ levels in *cup-5* defective nematodes are rebalanced (Figure 1B and 1C).

**Figure 1.**
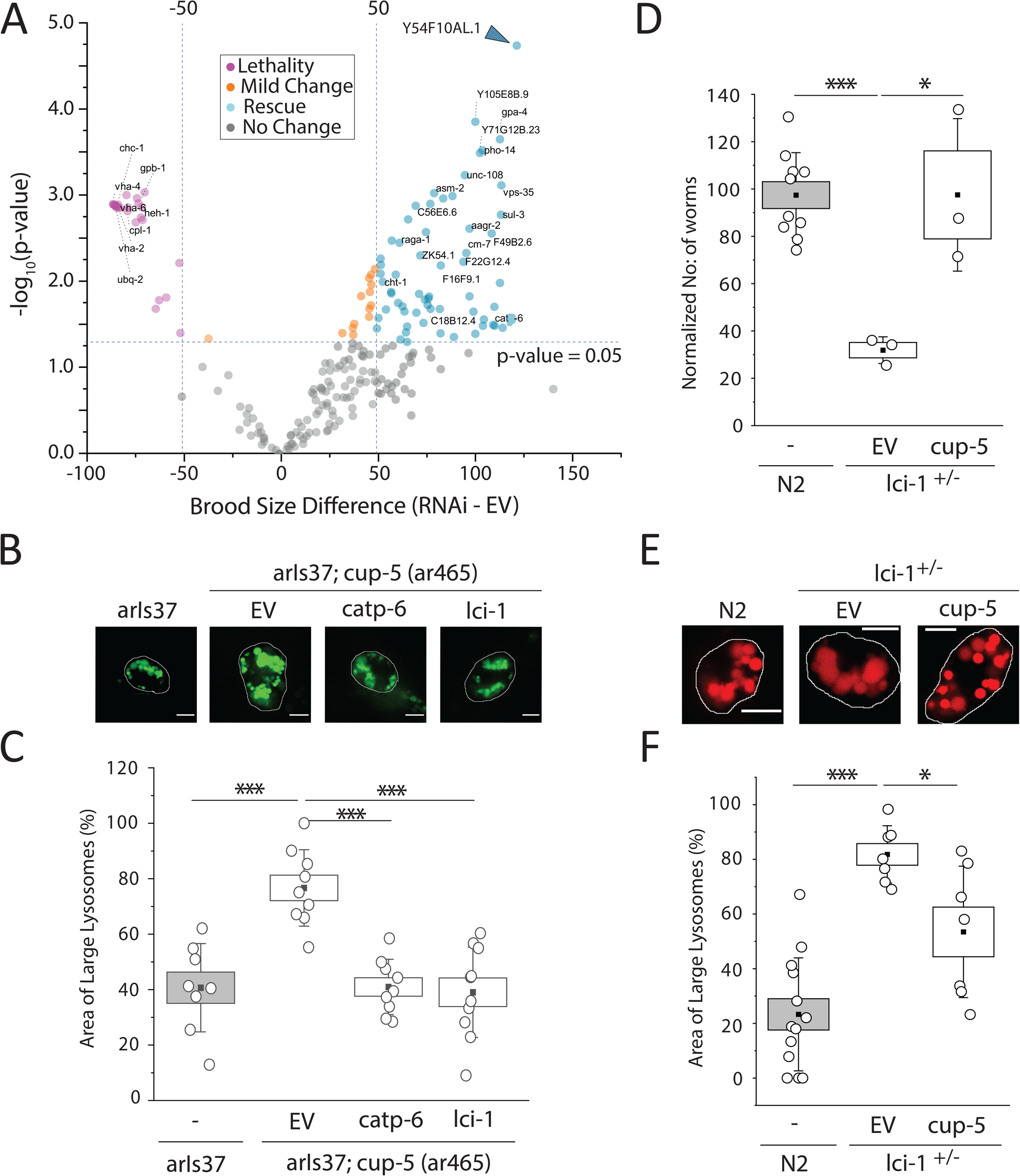
A phenotypic *sc*reen in *C. elegans* identifies *lci-1* as facilitating lysosomal Ca^2+^ import. (A) Survival of *cup-5*^+/-^ worms following RNAi knockdown of the 228 lysosomal genes, compared to worm survival following knockdown with an empty vector. Genes increasing survival (rescue) have a brood size change >50 and p-value <0.05, while genes decreasing survival (lethality) have brood size change <-50 and p-value <0.05. (B) Representative fluorescence images of lysosomes in coelomocytes of *arIs37* and *arIs37; cup-5(ar465)* worms following on RNAi knockdown of indicated proteins. (C) Percentage area occupied by enlarged lysosomes in the indicated genetic background. (n>10 cells, >75 lysosomes). (D) Number of N2 or *lci-1*^+/-^ progeny following RNAi knockdown of *cup-5*. (E) Representative fluorescence images of lysosomes in coelomocytes of *N2* and *lci-1^+/-^* worms in the indicated genetic background. Lysosomes are labeled with Alexa647 duplex DNA. (F) Percentage area occupied by enlarged lysosomes in the indicated genetic background. (n>5 cells, >50 lysosomes). *EV*, empty vector; *ncx*, Na^+^/Ca^2+^ exchanger; *clh*-*6*, Cl^-^ channel protein; *catp-6*, Ca^2+^-transporting ATPase; *lci-1*, lysosomal Ca^2+^/H^+^ exchanger. Scale bar, 5 μm. All experiments performed in triplicate. Boxes and bars represent the s.e.m. and standard deviation, respectively. *p<0.05; **p<0.01; ***p<0.001 (one-way ANOVA with Tukey post hoc test).

We then tested whether the relationship between *lci-1* and *cup-5* is commutative by seeing whether *cup-5* knockdown rescues *lci-1^-/-^*phenotypes. Homozygous *lci-1* knockout worms are embryonic lethal, *lci-1^+/-^* worms show small brood sizes and their coelomocytes have swollen lysosomes (Figure 1D-F and S1E-G). Knocking down *cup-5* in *lci-1^+/-^* worms rescues lethality, brood sizes (Figure 1D and S1H) and restores normal lysosome morphology (Figure 1E and 1F). Thus, *lci-1* acts in opposition to *cup-5*, likely facilitating lysosomal Ca^2+^ accumulation directly or indirectly.

### Human LCI facilitates lysosomal Ca^2+^ accumulation in nematodes

The gene *lci-1* is evolutionarily conserved from yeast (Gdt1) to humans (TMEM165)^21^. The human homolog, hereafter denoted LCI, is implicated in selected congenital disorders of glycosylation (CDG). While LCI is predominantly Golgi-resident, a significant fraction is also present in lysosomes^22, 23^. Missense mutations in human TMEM165 lead to it favoring either Golgi localization (compound heterozygous G304R and R126C) or lysosomal localization (homozygous R126C/H)^24^. In humans, loss of TMEM165 localization in either organelle causes growth and psychomotor retardation, muscular weakness, severe dwarfism, and death in infancy^23, 25^. Interestingly, R126 lies within a putative YNRL lysosomal-targeting motif (Figure 2A)^24^. In our hands too, LCI-EGFP is Golgi-resident, the R126C or R126H variants are predominantly in lysosomes, and the G304R variant is predominantly in the Golgi (Figure 2B-C and S2A-B and Note S1). A DsRed fusion of WT-LCI as well as endogenous LCI were clearly present on lysosomal membranes (Figure 2D-E and S2C)^26^, reaffirming that in addition to the Golgi, a small, yet significant, fraction resides in lysosomes.

**Figure 2.**
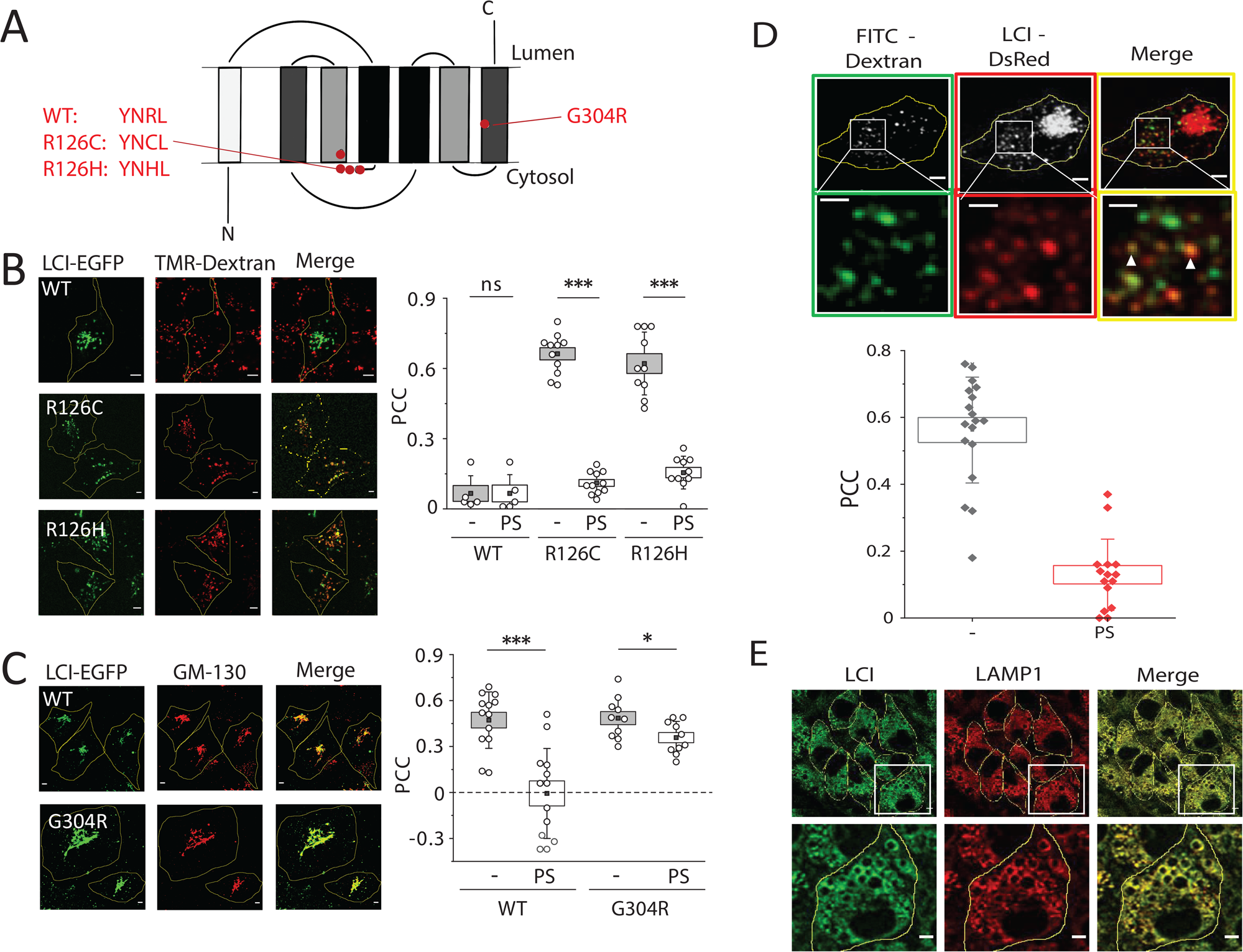
Human LCI partially localizes to lysosomal membranes. (A) Schematic of LCI topology showing internal symmetry of transmembrane helices and location of key mutations. (B) Left, representative fluorescence images of COS-7 cells transiently expressing the indicated variant of LCI-EGFP (green) and TMR-Dextran-labeled lysosomes (red). Right, Pearson correlation coefficient (PCC) of LCI and TMR-dextran before and after pixel shift (PS). n>10 cells (C) Left, representative fluorescence images of COS-7 cells transfected with the indicated variant of LCI-EGFP (green) and immunostained for GM130 (red). Right, PCC of LCI and GM130 before and after pixel shift (PS). n>10 cells. (D) Top, representative fluorescence images of COS-7 cells transiently expressing LCI-DsRed (red) and labeled with FITC-dextran (green). White arrows indicate LCI-positive lysosomes in inset merge image. Bottom, PCC of colocalization between LCI and FITC-dextran before and after pixel shift (PS). n>10 cells (E) Representative immunofluorescence images showing endogenous localization of LCI in HeLa cells pre-treated with vacuolin-1 to swell lysosomes. Cells were stained with antibodies to LCI (green) and LAMP1 (red) after glyoxal fixation to maintain lysosome structure. Scale bar 5 µm. Inset scale bar 2 µm. Boxes and bars represent the s.e.m. and standard deviation, respectively. I-shaped box represents the mean ± s.e.m. ns, not significant (p>0.05); *p<0.05; **p<0.01; ***p<0.001 (one-way ANOVA with Tukey post hoc test).

We then tested whether human LCI can rescue *lci-1* worm phenotypes, by extrachromosomally expressing either lysosome-favoring (R126C) or Golgi-favoring (G304R) LCI variants in these worms (Figure S2D). Only WT LCI and R126C LCI rescued the brood size and lysosome morphology defects of *lci-1*^+/-^ worms (Figure 3A-B and S2E-F). This indicates that restoring functionality in the Golgi alone is insufficient to rescue *lci-1* phenotypes and that the lysosomal fraction of LCI is physiologically significant.

**Figure 3.**
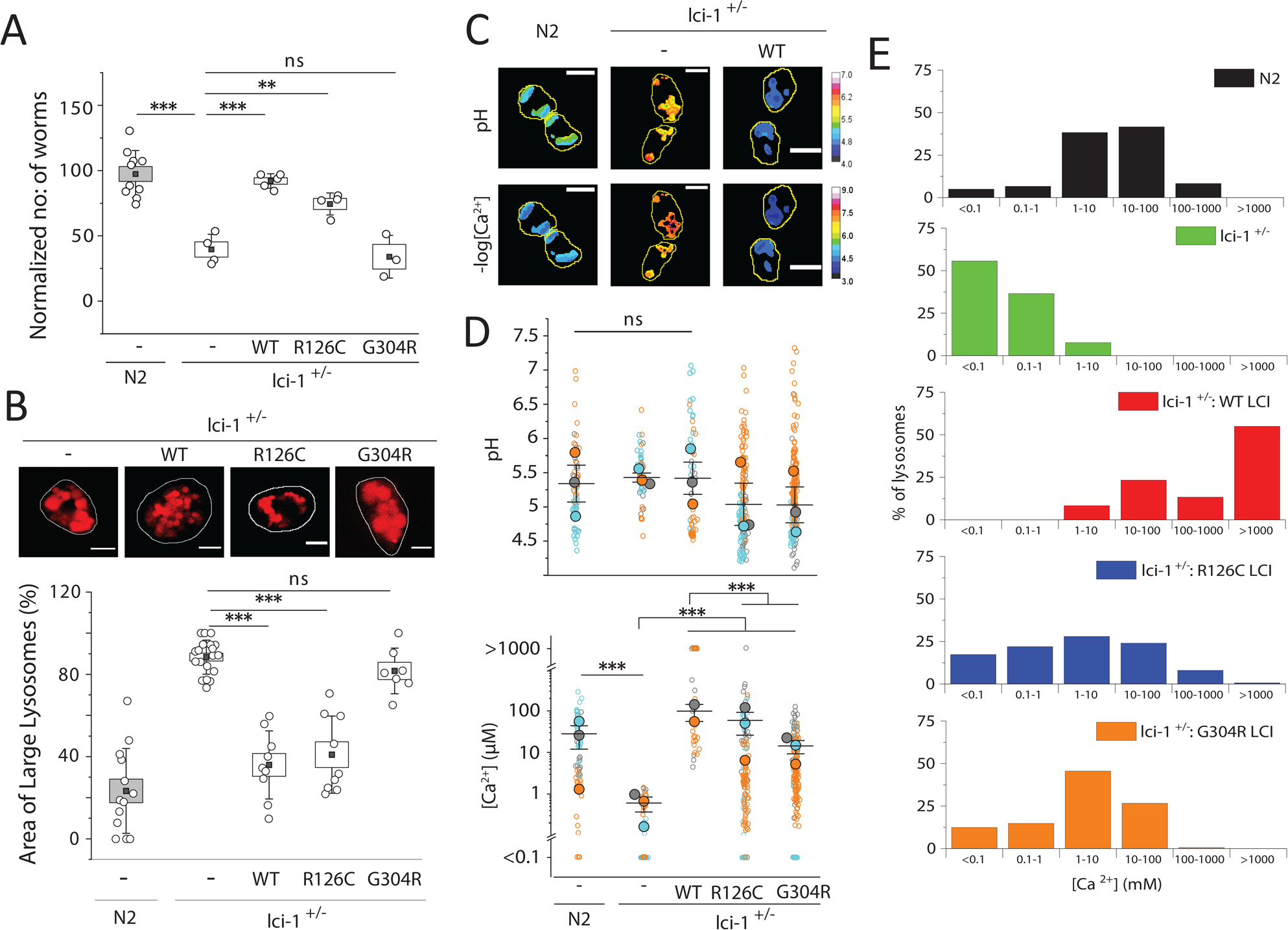
Human LCI rescues lysosomal Ca^2+^ import defects. (A) Number of progeny of *lci-1*^+/-^ worms extrachromosomally expressing the indicated human LCI variant (n=3 replicates). (B) Top, representative fluorescence images of lysosomes in coelomocytes of worms in the indicated genetic background. Lysosomes are labelled with Alexa647 duplex DNA. Bottom, percentage of area occupied by enlarged lysosomes in worms of the indicated genetic background. n>5 cells, >50 lysosomes. (C) Representative pH and –log([Ca^2+^]) maps in *CalipHluor2.0*-labeled lysosomes in coelomocytes in indicated genetic backgrounds (D) Distribution of lysosomal pH (top) and lysosomal Ca^2+^ (bottom) in the indicated genetic backgrounds. Data represents the means (closed blue, orange, and gray circles) of 3 independent experiments (open blue, orange, and gray circles). Lysosomes with Ca^2+^ levels below O/R_min_ or above O/R_max_ of *CalipHluor2.0* are plotted as <0.1µM and >1000µM, respectively. (E) Distribution of lysosomes with the indicated Ca^2+^ concentration measured using *CalipHluor2.0* in the indicated genetic backgrounds. Scale bar 5 µm. Boxes and bars represent the s.e.m. and standard deviation, respectively. I-shaped box represents the mean ± s.e.m. ns, not significant (p>0.05); *p<0.05; **p<0.01; ***p<0.001 (one-way ANOVA with Tukey post hoc test).

To understand why LCI rescues lysosome dysfunction in *lci-1* mutants, we measured pH and Ca^2+^ levels at single lysosome resolution, using a previously described DNA-based, pH-correctable Ca^2+^ reporter, *CalipHluor 2.0* (Figure S2G-H and Methods)^17, 27^. While lysosomal pH in *lci-1*^+/-^ worms did not vary significantly from those in N2 worms, lysosomal Ca^2+^ levels are ∼100-fold lower (Figure 3C-D and Note S2). Extrachromosomal expression of either WT LCI or the R126C LCI fully restores lysosomal Ca^2+^ levels, while G304R-LCI partially restored it (Figure 3D-E and S2I). These results indicate that the heterologous expression of human LCI facilitates Ca^2+^ accumulation in nematode lysosomes thereby restoring lysosome function and rescuing lethality.

### Mutations of putative Ca^2+^ binding sites impair LCI activity

Since TMEM165 was previously assigned as a Ca^2+^/H^+^ exchanger^21, 28^, we tested whether it compensates the well-known *S. cerevisiae* CAX gene, vcx1, which imports excess cytosolic Ca^2+^ into the vacuole in exchange for vacuolar H^+^[29,30]. Despite its lack of homology with CAX families or other cation/Ca^2+^ (CaCA) transporters, we looked for shared structural features between LCI and vcx1. LCI and its orthologs belong to the UPF0016 family of membrane proteins of unknown function. UPF0016 family proteins contain two EXGDK/R motifs flanked by two hydrophobic regions (Figure 4A-B)^23^. The acidic residues around the EXGDK/R motifs are posited to be involved in cation recognition^24^ and previous studies suggest that TMEM165 conducts Ca^2+^ current^21, 31, 32^. The structure of vcx1 too, revealed a cytosolic loop rich in acidic amino acids that coordinates Ca^2+^ ions which are transported into the vacuole via a conformational change induced by the transmembrane H^+^ gradient^29^. UPF0016 family proteins show two-fold antiparallel symmetry, just like vcx1 and other CaCA transporters (Tables S2 and S3)^21, 33^. LCI has only 7 predicted TM domains unlike canonical CAX transporters that have ∼14. However, it has regions with high homology to the acidic loop and to the helices in vcx1 that bind Ca^2+^ (Figure S3A). These regions are highly conserved within the UPF0016 family and are adjacent to the EXGDK/R motif (Figure 4B and S3B). Further, a homozygous missense E108G mutation in the EXGDK/R motif in TMEM165 leads to CDG Type II in humans^34^. A homology model of LCI based on vcx1 and other transmembrane proteins shows the putative cation-binding region lining a pore with direct access to the acidic cytosolic loop and the lysosome-targeting YNRL motif on the same side as the acidic helix (Figure S3C and Table S4 and Note S3). Based on these similarities, we tested whether LCI activity is affected by mutating the putative cation-binding regions and in the acidic loop.

**Figure 4.**
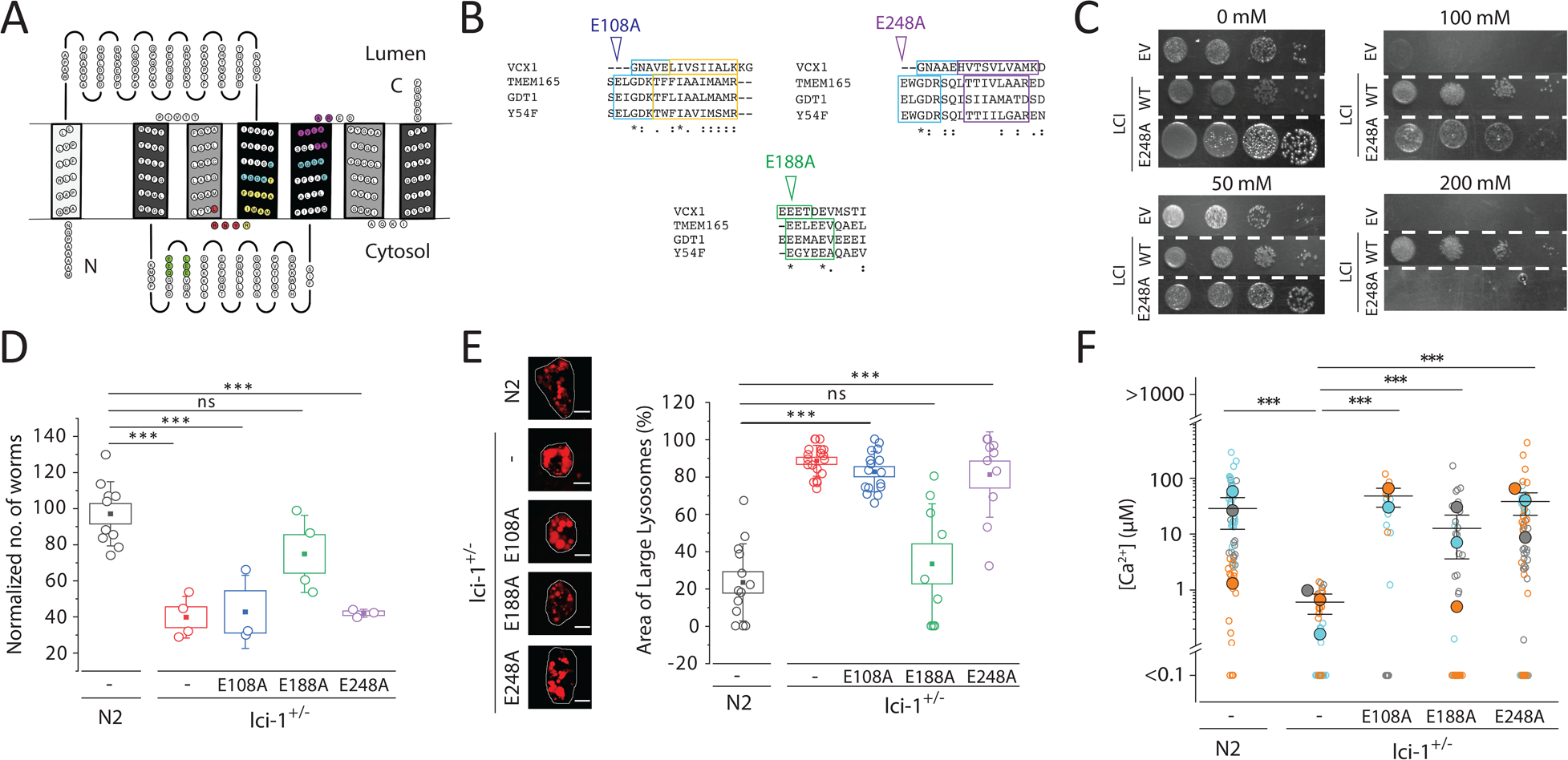
Homology with vcx1 reveals potential Ca^2+^-binding sites in LCI. (A) Topology of LCI created using TOPO2. Grayscale shows internal symmetry of transmembrane segments. Ca^2+^ binding regions (light blue), homologous regions adjoining the Ca^2+^ binding regions (yellow and purple), acidic helices (green), and lysosome/vacuole-targeting motifs (red) are highlighted. (B) Alignment of the Ca^2+^ binding sites (top and bottom) and acidic helix (middle) of vcx1 (*S. cerevisiae*), TMEM165/LCI (*H. sapiens*), *Y54F10AL.1*/*lci-1* (*C. elegans*), and gdt1 (*S. cerevisiae*) using Clustal Omega. Asterisks, colons and periods indicate full, strong and weak conservation, respectively. (C) Growth of K665 transformants integrating either an empty vector (EV) or the indicated variant of human LCI, after 2 days at 30°C on YPD plates supplemented with the indicated concentration of CaCl_2_. Columns indicate 10-fold dilutions from left-to-right. (D) Number of progeny of of *lci-1^+/-^* worms extrachromosomally expressing the indicated human LCI variant (n=3 replicates). (E) Left, representative fluorescence images of lysosomes in coelomocytes in worms of the indicated genetic background. Lysosomes are labelled with Alexa647 duplex DNA. Right, percentage of area occupied by enlarged lysosomes in the indicated genetic background. N>5 cells, >50 lysosomes. (F) Distribution of lysosomal Ca^2+^ in the indicated genetic backgrounds. Data represents the means (closed blue, orange, and gray circles) of 3 independent experiments (open blue, orange, and gray circles). Lysosomes with Ca^2+^ levels below O/R_min_ or above O/R_max_ of *CalipHluor2.0* are plotted as <0.1 µM and >1000 µM, respectively. Scale bar 5 µm. Boxes and bars represent the s.e.m. and standard deviation, respectively. ns, not significant (p>0.05); *p<0.05; **p<0.01; ***p<0.001 (one-way ANOVA with Tukey post hoc test).

We exogenously expressed LCI in an *S. cerevisiae* strain, denoted K665, that lacks both the vacuolar Ca^2+^ importers, pmc1 and vcx1^35^. High extracellular Ca^2+^ is toxic to K665 and exogenously expressing *A. thaliana* CAX genes reverses this lethality^35^. Exogenously expressing WT LCI in K665 reverses its Ca^2+^ sensitivity (Figure 4C and S3D-G and Note S4), while an E248A mutation in a putative cation-binding site of LCI, could not (Figure 4C). Extrachromosomal expression of E108A or E248A LCI variants, with mutations in either putative cation-binding site of LCI, also failed to rescue the brood size and lysosome size defects of *lci-1*^+/-^ worms (Figure 4D-E and S4A-B). This indicates that these mutations likely inactivate LCI. An E188A mutation in the acidic loop led to partial LCI activity in these assays (Figure 4D-E and S4A-B). None of these point mutations alter protein expression levels or lysosomal localization (Figure S4C-G) indicating that the phenotypic differences are due to protein activity. Expression of all mutants in worms only partially rescued lysosomal Ca^2+^ levels (Figure 4F and S4H-J and Note S2). Cumulatively, the results indicate that the EXGDK/R motif is vital for LCI’s ability to elevate lysosomal Ca^2+^, as seen previously for vcx1^31^. This suggests functional similarity between LCI and vcx1, despite their low overall structural homology.

### LCI facilitates Ca^2+^ entry into human lysosomes pH-dependently

We then tested whether LCI facilitated Ca^2+^ entry into lysosomes of human cells. First, we elevated cytosolic Ca^2+^ with ATP in HeLa cells expressing LCI, tracked the cytosolic Ca^2+^ spike and its subsequent decrease by following Fura red fluorescence with time (Figure 5A)^36^. We also mapped lysosomal pH under identical conditions by imaging FITC-dextran labeled lysosomes in whole cells (Figure 5A). An overlay of both traces reveals that lysosomal pH increases during the decay period of the cytosolic Ca^2+^ spike. Our findings are consistent with that of others, which led to them positing the existence of a pH-dependent lysosomal Ca^2+^ entry mechanism, such as Ca^2+^/H^+^ exchange^14, 16^. We therefore measured Ca^2+^ and pH in single lysosomes of HeLa cells where LCI is either knocked out or overexpressed (Figure S5A and S5B), using a *CalipHluor* variant called *CalipHluor^mLy^*, suited to probing mammalian lysosomes (Figure S5C and S5D)^17^. We found that on average, LCI deletion decreases lysosomal pH by ∼0.5 units, and decreases lysosomal Ca^2+^ by 5-fold, which suggests a Ca^2+^/H^+^ exchange model of LCI (Figure 5B-E and S5E).

**Figure 5.**
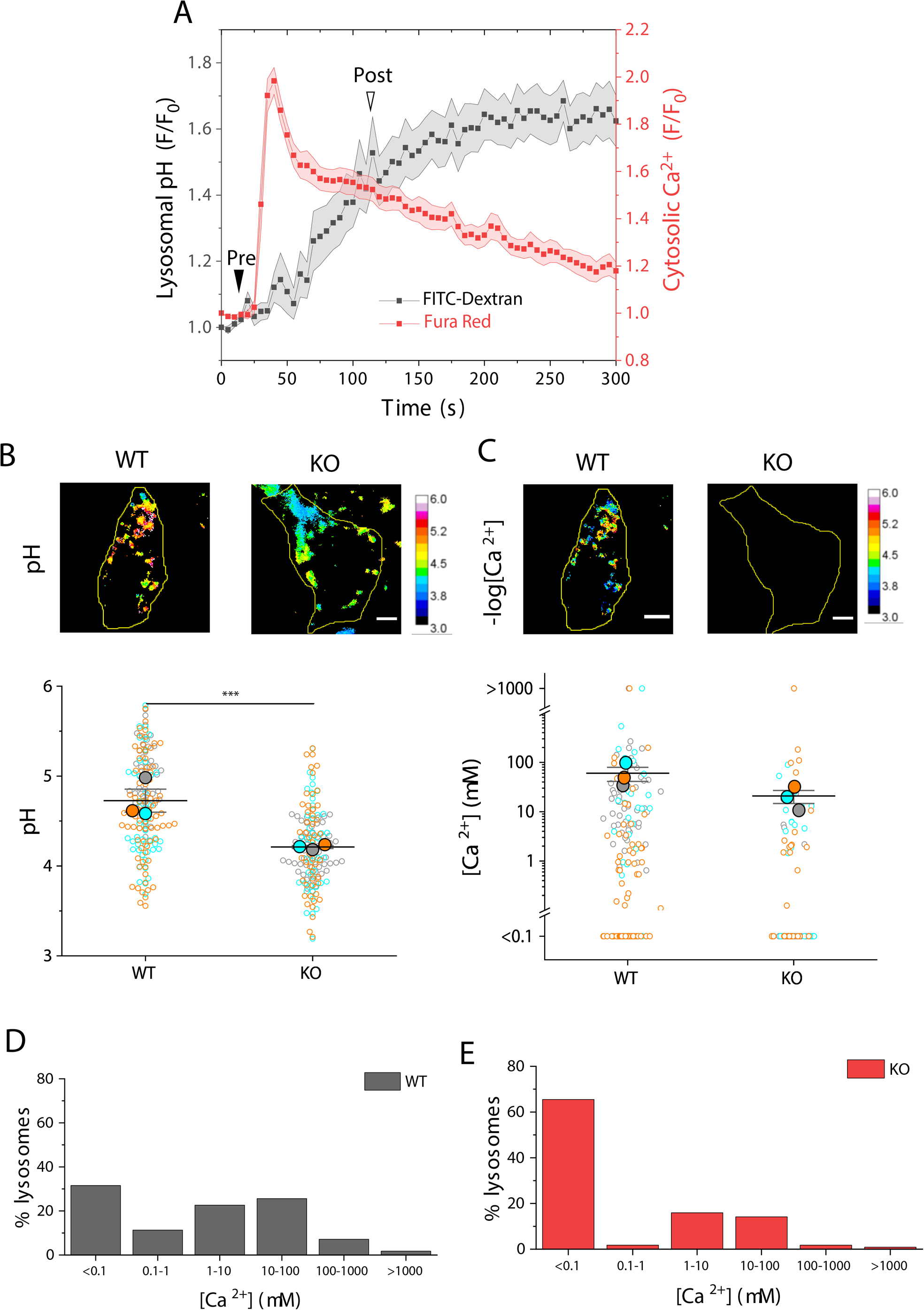
Population-averaged lysosomal pH and Ca^2+^ measurements indicate Ca^2+^/H^+^ exchange. (A) Overlay of the cytosolic Ca^2+^ recovery and lysosomal pH spike, given by the 440/488 ratio of Fura Red and the 488/440 ratio of FITC-dextran, respectively, in WT cells. Arrowheads indicate imaging time points before ATP addition (Pre, filled arrow) and after ATP addition (Post, unfilled arrow) in Figure 6. (B) Top, representative pH maps of lysosomes from WT and LCI KO HeLa cells using *CalipHluor^mLy^*. Bottom, pH of individual lysosomes (open circles) from three different experiments (closed circles). (C) Top, representative –log([Ca^2+^]) maps of lysosomes from WT and LCI KO HeLa cells using *CalipHluor^mLy^*. Bottom, [Ca^2+^] of individual lysosomes (open circles) from three different experiments (closed circles). Lysosomes below O/R_min_ or above O/R_max_ of *CalipHluor^mLy^* are shown as <0.1µm Ca^2+^ and >1000µm Ca^2+^, respectively. (D,E) Distribution of lysosomes with the indicated Ca^2+^ concentration measured using *CalipHluor2.0* in the indicated genetic backgrounds. Scale bar 5 µm. ***p<0.001 (one-way ANOVA with Tukey post hoc test).

To directly visualize Ca^2+^/H^+^ exchange in single lysosomes due to LCI activity, we simultaneously measured the pH and Ca^2+^ of single lysosomes at two different times (Figure 5A). One timepoint was in single lysosomes prior to cytosolic Ca^2+^ elevation. The other was 2 minutes after ATP addition corresponding to the Ca^2+^ spike, when the lysosomal Ca^2+^ entry mechanism, if any, is expected to be active. Our findings enforced a reconsideration of the Ca^2+^/H^+^ exchange model. We observed that after the cytosolic Ca^2+^ spike, pH increases only in 20% of lysosomes, decreases in 50% of lysosomes, and was unchanged in the remaining 30% (Figure 6A), revealing the limitations of population-averaged lysosome measurements and the pH range quantifiable by FITC-dextran. Yet, Ca^2+^ increases in about 72% of lysosomes (Figure 6B). When we performed the same experiments in LCI KO HeLa cells, the population of lysosomes where pH decreased in response to high cytosolic Ca^2+^, was lost (Figure 6C and 6D).

**Figure 6.**
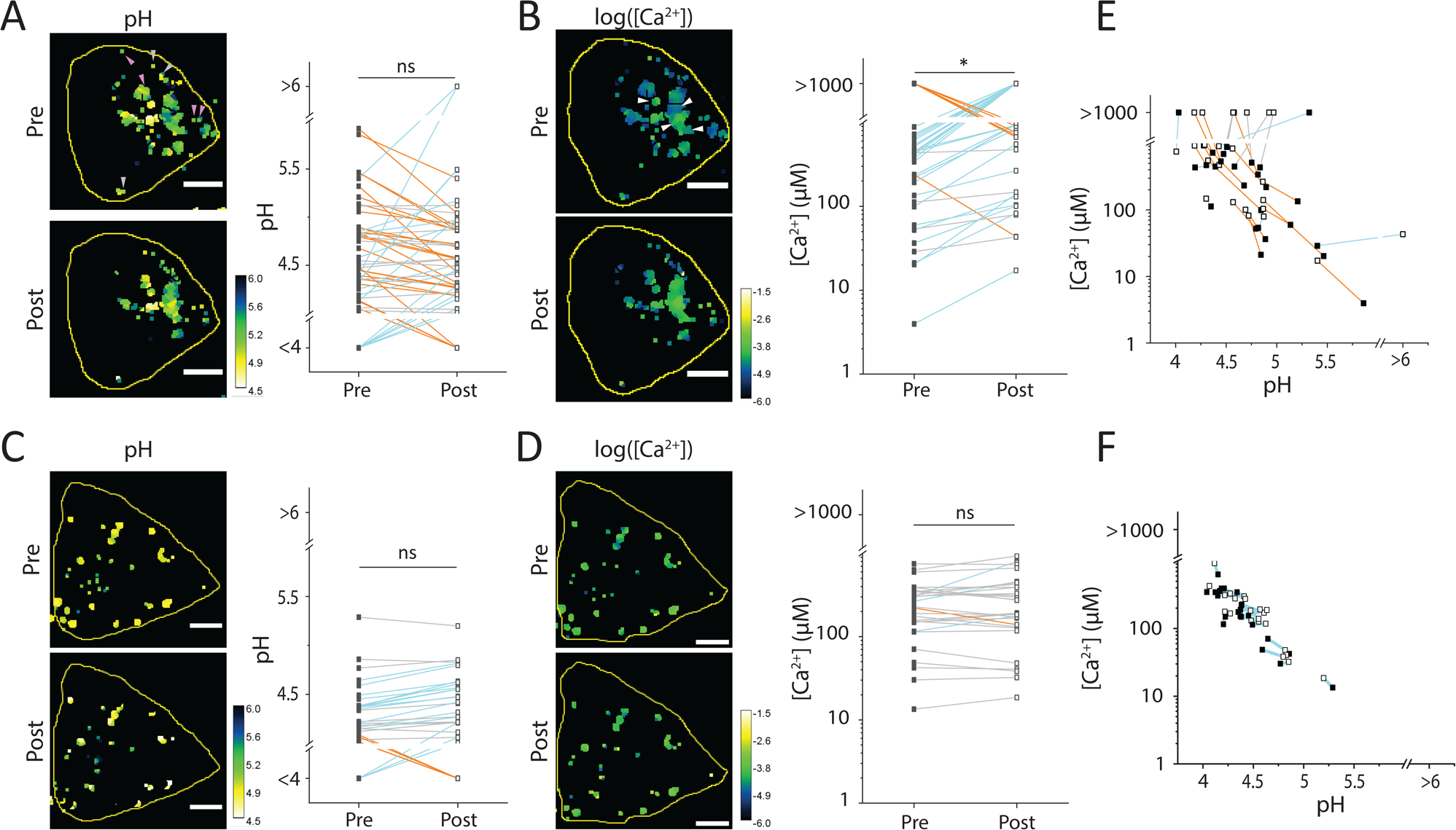
Two-ion mapping at single lysosome resolution indicates H^+^-dependent Ca^2+^ import. (A) Left, representative pH maps of lysosomes from WT HeLa cells using *CalipHluor^mLy^* before and 2 minutes after 100 µM ATP addition. Arrowheads indicate lysosomes where pH decreases. Right, pH change of individual lysosomes of WT HeLa cells following addition of ATP. Increasing, decreasing, and unchanged pH are indicated by blue, orange, and gray lines, respectively. (B) Left, representative log([Ca^2+^]) maps of lysosomes from WT HeLa cells using *CalipHluor^mLy^* before and 2 minutes after 100 µM ATP addition. Arrowheads indicate lysosomes where [Ca^2+^] increases. Right, [Ca^2+^] changes in individual lysosomes of WT HeLa cells following ATP addition. Increasing, decreasing, and unchanged [Ca^2+^] are indicated by blue, orange, and gray lines, respectively. (C) Left, representative pH maps of lysosomes from LCI KO HeLa cells using *CalipHluor^mLy^* before and 2 minutes after addition of 100 µM ATP. Right, pH change of individual lysosomes of LCI KO HeLa cells following addition of ATP. Increasing, decreasing, and unchanged pH are indicated by blue, orange, and gray lines, respectively. (D) Left, representative log([Ca^2+^]) maps of lysosomes from LCI KO HeLa cells using *CalipHluor^mLy^* before and 2 minutes after addition of 100 µM ATP. Right, [Ca^2+^] change of individual lysosomes of LCI KO HeLa cells following addition of ATP. Increasing, decreasing, and unchanged [Ca^2+^] are indicated by blue, orange, and gray lines, respectively. (E) 2-IM maps of pH and [Ca^2+^] changes in WT HeLa cells before (closed square) and after (open square) adding ATP. Orange lines indicate lysosomes whose pH decreases and Ca^2+^ increases, cyan lines indicate lysosomes whose pH increases and Ca^2+^ increases, and gray lines indicates lysosomes whose Ca^2+^ decreases. (F) 2-IM maps of pH and [Ca^2+^] changes in LCI KO HeLa cells before (closed square) and after (open square) adding ATP. Orange lines indicate lysosomes whose pH decreases and Ca^2+^ increases, cyan lines indicate lysosomes whose pH increases and Ca^2+^ increases, and gray lines indicates lysosomes whose Ca^2+^ decreases. Scale bar 5 µm. ns, not significant (p>0.05); *p<0.05 (one-way ANOVA with Tukey post hoc test).

To better reveal the missing lysosome population when LCI was knocked out, we plot the pH of every lysosome versus its Ca^2+^ level to give a scatter plot, called a two-ion measurement plot, or 2-IM plot. 2-IM plots have been previously used to resolve lysosome populations in live cells^37^. The 2-IM plots of HeLa cells expressing (Figure 6E) and lacking LCI (Figure 6F) show lines connecting the same lysosome pre and post ATP addition. For clarity, orange lines show those lysosomes where pH decreases but Ca^2+^ increases, while cyan lines show lysosomes where both pH and Ca^2+^ increase, post ATP addition. Note that Ca^2+^/H^+^ exchange is expected to favor the cyan population. Yet, after the cytosolic Ca^2+^ spike, nearly 75% of lysosomes that showed Ca^2+^ increase, were also more acidic. This is the opposite of what is expected if TMEM165 was a Ca^2+^/H^+^ exchanger as previously posited.

There are two possible explanations for the concurrent increase in Ca^2+^ and H^+^ levels in single lysosomes. One is an active mechanism such as pH-dependent Ca^2+^ transport^14, 38^. Alternatively, lumenal Ca^2+^ could get passively elevated because higher acidity protonates Ca^2+^-binding lysosomal proteins, making them release their bound Ca^2+^. The contribution of the passive increase in free Ca^2+^ is reflected in the negative slope of both 2-IM plots - pre and post Ca^2+^ spike - in LCI KO cells (Figure S5F and S5G). When LCI is absent, the contribution of the passive mechanism is similar for both low and high cytosolic Ca^2+^ since protein content in single lysosomes is not expected to differ between both time-points. The passive mechanism is also evident in HeLa cells expressing LCI, but only at low cytosolic Ca^2+^ (Figure S5H). However, at high cytosolic Ca^2+^, the slope increases, indicating much stronger coupling between H^+^ and Ca^2+^ levels in lysosomes when LCI is present (Figure S5I). This reveals that LCI is responsible for an additional Ca^2+^ entry pathway in lysosomes that depends on protons.

### Electrophysiology establishes LCI as a pH-dependent Ca^2+^ importer

To test whether LCI directly transported Ca^2+^ across cellular membranes we performed whole-cell patch-clamp electrophysiology using NMDG-MSA buffers, where Ca^2+^ was the only transportable ion. As LCI overexpression leads to adequate cell surface expression (Figure S6A), we mimicked, at the plasma membrane, LCI activity across the lysosome membrane by using NMDG-MSA bath and pipette buffers that mimicked lysosomal and cytosolic ionic environments respectively (Figure 7A and S6B)^28^. In the absence of a transmembrane pH gradient, HeLa cells overexpressing LCI showed significantly higher outward currents than mock-transfected HeLa cells indicating the movement of Ca^2+^ from the cytosol into the bath/lysosome even at low cytosolic Ca^2+^ (Figure 7B and S6C). To explicitly test whether LCI transports Ca^2+^, we progressively reduced Ca^2+^ in the cytosol/pipette buffer and recorded currents as a function of the Ca^2+^ gradient alone (Figure 7C). With only an inward Ca^2+^ gradient, we observed high current density of 30 pA/pF at 100 mV with the direction of current matching the flow of Ca^2+^ down the gradient. When cytosolic Ca^2+^ is reduced to trace levels, where pipette/cytosolic buffer contains only EGTA, two things change: the outward current decreases three-fold, and the reversal potential shifts towards slightly more positive values (Figure 7C-D and S6D). Thus, LCI directly transports Ca^2+^ across cellular membranes.

**Figure 7.**
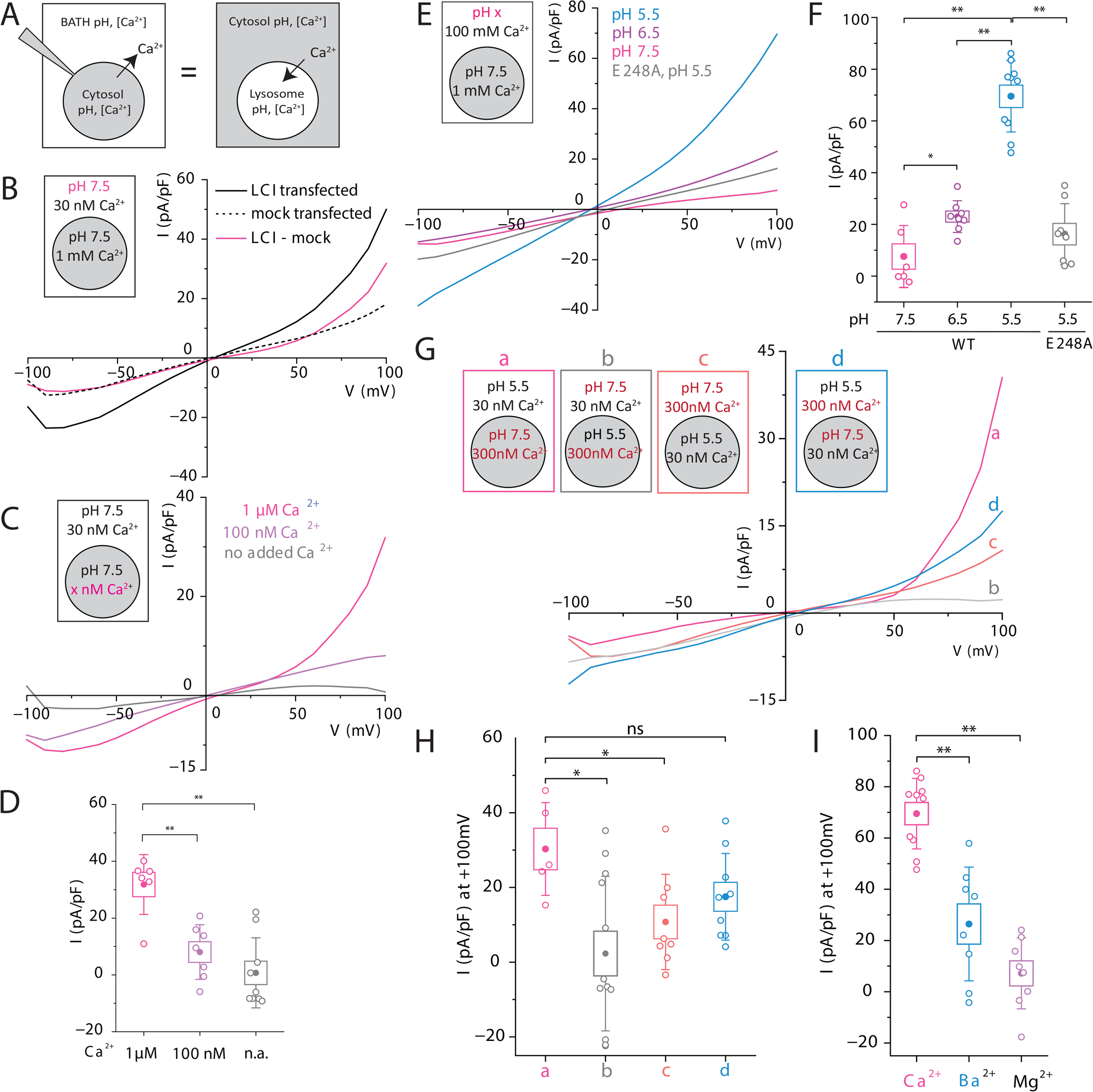
LCI transports Ca^2+^ unidirectionally and pH-dependently into lysosomes. (A) Schematic of whole-cell electrophysiology used to record LCI currents. Bath/extracellular solution is equivalent to the lysosomal lumen, and the pipette solution is equivalent to the cytosol. Outward (positive) current represents Ca^2+^ transport from the cytosol to the bath/lysosome lumen. (B) Average current densities of mock-transfected HeLa cells (dashed line), LCI-transfected HeLa cells (solid black line) and mock-subtracted LCI current density (solid magenta line). (C) Average current densities of mock-subtracted LCI current in HeLa cells under the indicated conditions, with no added Ca^2+^ (gray), 100 nM Ca^2+^ (mauve), or 1 µM Ca^2+^ (magenta). (D) Current density at +100mV for the indicated conditions in (c). (E) Average current density of mock-subtracted E248A (gray) or WT LCI current in HeLa cells under the indicated conditions, with an extracellular pH of 7.5 (magenta), 6.5 (mauve), or 5.5 (blue). (F) Current density at +100mV for the indicated conditions in (e). (G) Average current density of mock-subtracted LCI current in HeLa cells for the indicated conditions, with either 30 nM or 300 nM Ca^2+^ and pH 5.5 or 7.5 in the cytosol and bath. (H) Current densities at +100mV for the indicated cell conditions from (g). (I) Average current density at +100mV for conditions indicated in Figure S6G. Boxes and bars represent the s.e.m. and standard deviation, respectively. ns, not significant (p>0.05); *p<0.05; **p<0.01; ***p<0.001 (one-way ANOVA with Tukey post hoc test).

We then studied the pH-dependence of Ca^2+^ transport by LCI. We recorded currents under increasing transmembrane pH gradients, by increasing the acidity of the bath/lysosomal buffer, while keeping the inward Ca^2+^ gradient constant (Figure 7E). We found that the outward currents conducted by LCI increased with increasing acidity of the bath/lysosomal buffer, after subtracting mock-transfected currents under identical conditions (Figure 7E and 7F). Thus, the transport of Ca^2+^ by LCI increases with increasing acidity of the bath/lysosomal buffer. Overexpressing LCI with an alanine mutation in its putative cation-binding site (E248A) reduced the outward current nearly 3-4 fold, even at high transmembrane pH gradients, indicating that this mutation heavily compromises Ca^2+^ transport (Figure 7E and 7F).

TMEM165 was previously postulated to be a Ca^2+^/H^+^ exchanger (CAX) because it feebly resembled the vacuolar CAX, vcx-1. However, electrophysiology as a function of pH reveals reversal potential shifts of <10 mV in the negative direction for a 100-fold difference in transmembrane H^+^ (ΔpH) which is miniscule for a CAX. (Figure S6E). Even a 10-fold change in [ΔH^+^] should shift the reversal potential by ∼50-180 mV in the positive direction for reasonable stoichiometries (Table S5). Thus, even though protons enhance Ca^2+^ transport, H^+^ transport *per se* by LCI is insignificant (Figure 7E).

Exchangers can switch their ion transport directionality depending on the transmembrane ion gradient. So, we tested the direction of Ca^2+^ transport by LCI at high (300 nM) or low (30 nM) cytosolic Ca^2+^; conditions expected to mimic the direction of Ca^2+^ flow across the lysosomal membrane in stimulated or resting cells, respectively (Figure 7G(a,d)). Interestingly, regardless of the direction of the transmembrane Ca^2+^ gradient LCI conducts predominantly outward Ca^2+^ current (Figure 7G and 7H). Since lysosome lumens are normally acidic, these results indicate that LCI imports Ca^2+^ into lysosomes regardless of an outward transmembrane Ca^2+^ gradient as in resting cells or an inward one as in excited cells or at an ER-Lysosome contact site. To try and reverse the direction of Ca^2+^ transport as seen in exchangers, we reversed the transmembrane pH and Ca^2+^ gradients and recorded currents (Figure 7G(b,c)). At acidic cytosolic pH, outward currents reduced ∼4-fold, while inward current changes were negligible (Figure 7H and S6F). Thus, LCI is only capable of transporting Ca^2+^ unidirectionally into the lysosome and is pH-activated only at the extracellular/lumenal face of the membrane. Further, the current amplitude through LCI as a function of divalent cations varies as Ca^2+^ > Ba^2+^ ∼ Mg^2+^ (Figure 7I and S6G). This behavior is unlike that of Ca^2+^ channels and more consistent with that of Ca^2+^ uniporters, such as MCU, that show a strong preference for Ca^2+^ [39–41].

Finally, we directly interrogated LCI current on isolated swollen lysosomes of COS7 cells, whose flatter morphology lends itself to lysosome patching. Both WT and R126C LCI-EGFP can be visualized on lysosomes of COS7 cells upon treatment with vacuolin-1, which swells lysosomes (Figure S7A and S7B)^42^. We used pipette buffers to mimic the lumenal acidity of either lysosomes or the Golgi (Figure S7C). Here too, lysosomes of cells expressing LCI show a large outwardly rectifying current (positive current into the lysosome) under either condition. At +100 mV, the current at lysosomal pH was ∼2.5 fold higher than at pH levels corresponding to the Golgi (Figure S7D and S7E). Cumulatively, our data is consistent with LCI being a proton activated, Ca^2+^ importer in lysosomes.

## Discussion

The high abundance of LCI (TMEM165) on the Golgi has led to detailed studies into its roles in protein glycosylation and its effects on human physiology^22, 32, 43–45^. While the most common pathophysiology of TMEM165 is linked to Golgi dysfunction, its significance in lysosomes has thus far been overlooked. For example, in a patient with heterozygous missense mutations (c.376C>T [p.R126C]) and (c.910G>A [p.G304R]), LCI is absent in lysosomes. Despite its presence on the Golgi, this patient manifested a less severe form of CDG-II, presenting with characteristic deformities that are also observed in lysosomal storage disorders such as fucosidosis and Gaucher’s disease^24^.

While LCI performs important Golgi-related roles, our data now show that it has an additional function, namely importing Ca^2+^ into lysosomes. LCI was previously posited to act like a Ca^2+^/H^+^ exchanger at the Golgi, based on its weak structural homology with vcx1. However, our studies reveal that LCI imports Ca^2+^ into organelle lumens by acting as a proton-activated Ca^2+^ importer. LCI is roughly half the size of a canonical CAX. We therefore suggest a model where LCI dimerizes to form a “broken CAX”, in which the proton-transport function is damaged, but Ca^2+^ can still be transported.

Lysosomal Ca^2+^ import is a key function in cells, and, while the existence of a pH-dependent Ca^2+^ import mechanism has long been known, the identity of the gene responsible was not. We now identify LCI as a pH-activated lysosomal Ca^2+^ importer in humans. Mutations in every known lysosomal Ca^2+^ channel disrupt lysosomal Ca^2+^ homeostasis and lead to diverse neurological disorders. Given that LCI can restore lysosomal Ca^2+^ channel phenotypes, it could provide new avenues to explore disorders arising from defective lysosomal Ca^2+^ channels.

## Supporting information

Key Resources Table

## Acknowledgements

We thank E. Perozo and J. Kuriyan for comments and discussions, P. Nandigrami and B. Roux for advice on homology modelling, K. Chakraborty and D. Mansour for technical assistance. We thank the integrated light microscopy facilities at the University of Chicago. Funding: This work was supported by the Women’s Board of the University of Chicago; FA9550-19-0003 from AFOSR, NIH grants R21NS114428, 1R01NS112139-01A1, and the Ono Pharma Foundation.

## Author Contributions

M.Z., A.S., and Y.K. conceptualized the project and initial experiments. M.Z., P.A., D.O, and J.S. performed experiments in worms. M.Z., K.H., and P.A. performed ion sensing experiments in cell lines. M.Z. and K.H. performed bioinformatics analyses. M.Z. and J.Z. performed experiments in yeast. S.M., A.S., and M.Z. performed electrophysiology experiments. M.Z., A.S., and S.M. analyzed data. M.Z., Y.K., and A.S. wrote the paper. All authors provided input on the manuscript.

## STAR Methods

### Resource Availability

#### Lead contact

Further information and requests for resources and reagents should be directed to and will be fulfilled by the lead authors Yamuna Krishnan and Anand Saminathan.

#### Materials availability

All reagents generated in this study are available from the lead author with a completed Material Transfer Agreement.

#### Data and code availability

There are not restrictions on data availability in this manuscript. Raw data files are available from the authors upon request.

### Experimental Model and Study Participant Details

#### Animal Models

Standard methods were used to maintain *C. elegans*^46^. The wild-type strain was the *C. elegans* isolate from Bristol, strain N2. Three strains used in the study were provided by the Caenorhabditis Genetics Center: *+/mT1 II; cup-5(ok1698)/mT1 [dpy-10(e128)] III*, a heterozygous knockout of *cup-5* balanced by *dpy-10* marked translocation; *arIs37[myo-3p::ssGFP+dpy-20(+)]I,* a transgenics strain expressing soluble GFP (ssGFP) in the body muscles which is secreted into the pseudocoelom and endocytosed by coelomocytes; and *arIs37[myo-3p::ssGFP+dpy-20(+)]Icup5(ar465)*, a transgenic strain with enlarged GFP-containing vesicles in coelomocytes due to a point mutation in *cup-5*. The strain with a heterozygous knockout of *lci-1* was generated and provided by D. Moerman (University of British Columbia) using CRISPR/Cas9 technology and verified by sequencing^47^. Heterozygous *lci-1*^+/-^ worms are marked by pharyngeal GFP, homozygous *lci-1*^+/+^ progeny are functionally wild-type but lack GFP, and homozygous *lci-1*^-/-^ progeny are embryonic lethal. The genotype of this worm is *Y54F10AL.1(gk5484[loxP + Pmyo-2::GFP::unc-54 3’ UTR + Prps-27::neoR::unc-54 3’ UTR + loxP])/+ III*. Transgenic strains expressing human LCI variants were generated by microinjecting of plasmid DNA into *lci-1^+/-^* gonads to produce extrachromosomal arrays. The injected plasmid contained an LCI variant with the promoter region of *lci-1* and the 3’ UTR of *unc-54*. Pharyngeal mCherry was used as an injection marker. The plasmid construction, worm injection, and verification by sequencing was performed by SunyBiotech (Fujian, China) using established protocols.

#### Cell lines

HeLa cells and COS7 cells were purchased from ATCC and cultured according to recommended guidelines. LCI KO HeLa cells were purchased from Creative Biogene. Cells were cultured in Dulbecco’s Modified Eagle’s Medium (DMEM) (Invitrogen Corporation, USA) containing 10% heat-inactivated fetal bovine serum (FBS) (Invitrogen Corporation, USA), 100 U/mL penicillin and 100 μg/mL streptomycin (Gibco), and maintained at 37°C under 5% CO_2_. Cells were passaged using 0.25% Trypsin-EDTA (Gibco) and plated at 50-60% confluency for transfection.

#### Yeast strains

The yeast strain K665 (*pmc::TRP1 vcx*Δ) was a kind gift from K. Hirschi (Baylor College of Medicine).

### Method Details

#### Chemicals and reagents

Source of chemicals and reagents are indicated in the Key resources table.

#### Plasmids and transfection in mammalian cells

For mammalian expression of LCI fused to EGFP, the cDNA of TMEM165 (Harvard Medical School Plasmid Database) was cloned into the pEGFP-N1 plasmid, which was obtained from M. Fransen (KU Leuven). Cloning was done using Gibson assembly techniques. For mammalian expression of LCI fused to DsRed, the DNA for DsRed (a kind gift from B. Dickinson at the University of Chicago) was cloned into the LCI-EGFP plasmid, replacing EGFP, using Gibson assembly techniques.

For mammalian expression of LCI disease mutants fused to EGFP, the above construct was subject to site-directed mutagenesis using the Q5 Site-Directed Mutagenesis Kit (NEB). The primers used are indicated in the Key Resources Table. Resulting plasmids were verified by sequencing. The GFP-Rab7 plasmid was a kind gift from Richard Pagano (Addgene plasmid #12605). HeLa and COS7 cells were transiently transfected with the respective plasmids using Lipofectamine 3000 (Thermo Fisher) according to manufacturer protocols. After incubation for 4 hours, the transfection medium was replaced with fresh DMEM. Imaging or electrophysiology experiments were performed on cells 48h following transfection.

#### RNAi and RT-PCR in *C. elegans*

Gene knockdown was performed using Ahringer Library-based RNAi methods^48^. The RNAi clones used were L4440 empty vector control, *catp-6* (W08D2.5, Ahringer Library)*, cup-5* (R13A5.1, Ahringer Library), *lci-1* (Y54F10AL.1, Ahringer Library)*, ncx-1* (Y113G7A.4, Ahringer Library)*, ncx-2* (C10G8.5, Ahringer Library)*, ncx-3* (ZC168.1)*, ncx-4* (F35C12.2), and *clh-6* (R07B7.1, Ahringer Library).

Bacteria from the Ahringer RNAi library (obtained as a kind gift from F. Ausubel, Massachusetts General Hospital) expressing double-stranded RNA against the relevant gene were fed to worms, and the relevant experiments were carried out in 1d-old adults of the F1 progeny. RNA knockdown and genetic background were confirmed by probing messenger RNA levels of the candidate gene, assayed by RT-PCR. Briefly, total RNA was isolated using the Trizol-chloroform method and 1 ug total RNA was converted to complementary DNA using oligo-dT primers and SuperScriptIV RT according to manufacturer protocols. Then, 5 µL of cDNA product was used to setup up a PCR using gene-specific primers indicated in the Key Resources Table. PCR products were separated on a 0.8% agarose-Tris base, acetic acid, and EDTA (TAE) gel.

To confirm expression of LCI-1 in the *S. cerevisiae* strain K665, RNA levels were probed using RT-PCR. gDNA was isolated using Monarch Genomic DNA Purification Kit (NEB) according to manufacturer protocols. Briefly, transformants were grown overnight in liquid SD-Leu media and cells were harvested by centrifugation for 1 minute at 12,000 x g. Cells were resuspended in lysis buffer with 10 µL zymolyase and 3 µL RNase A and incubated for 30 minutes at 37°C. Then, 10 µL Proteinase K and 100 µL Tissue Lysis Buffer were added and the mixture was vortexed thoroughly before incubating for 30 minutes at 56°C with shaking. Then, 400 µL of gDNA Binding Buffer was added, and the mixture was vortexed before transferring to a purification column in a collection tube. The column was centrifuged for 3 minutes at 1,000 x g and then 1 minute at 12,000 x g. Then, the column was washed twice with 500 µL of gDNA Wash Buffer before adding 50 µL pre-heated gDNA elution buffer. The column was incubated for 1 minute and centrifuged into a microforge tube for 1 minute at 12,000 x g. The purity and amount of DNA was determined using a NanoDrop, and 20ng of DNA was used to run a PCR using gene-specific primers indicated in the Key Resources Table. PCR products were separated on a 0.8% agarose-Tris base, acetic acid, and EDTA (TAE) gel.

#### Survival assay in *C. elegans*

The *N2*, *+/mT1 II; cup-5(ok1698)/mT1 [dpy-10(e128)] III,* and *Y54F10AL.1(gk5484[loxP + Pmyo-2::GFP::unc-54 3’ UTR + Prps-27::neoR::unc-54 3’ UTR + loxP])/+ III* nematode strains were used for this assay, which was performed as previously described^17^.

To screen lysosomal genes for calcium import activity, hundreds of L1-L2 worms of the indicated genotype were placed on fresh plates with OP50 and allowed to grow to L4 for 2 days. Five of these L4 worms were placed on plates containing bacterial strains L4440 with RNAi for the indicated genes. The worms were allowed to grow for 24 hours and lay eggs, after which the adult worms were removed from the plates. Eggs were allowed to hatch and grow into adults for 3 days. The worm plates were scanned twice on an Epson Perfection v850 Scanner in batches of 12 plates. The average number of worms on the plates between the two scans was calculated using the Fiji plug-in “WormAnalysisProgram.” The difference in brood size and statistical significance were calculated with respect to the empty vector.

#### Lysosome size assay in C. elegans

The *N2*, *arIs37[myo-3p::ssGFP+dpy-20(+)]I, arIs37[myo-3p::ssGFP+dpy-20(+)]Icup5(ar465),* and *Y54F10AL.1(gk5484[loxP + Pmyo-2::GFP::unc-54 3’ UTR + Prps-27::neoR::unc-54 3’ UTR + loxP])/+ III* nematode strains were used for this assay, which was performed as previously described^17^. Worms were imaged on a confocal microscope for either ssGFP-labeled coelomocyte lysosomes or a DNA duplex with Alexa647 (Key Resources Table).

#### Survival assay in *S. cerevisiae*

The yeast integrating plasmid YIplac128 was a kind gift from B. Glick (University of Chicago). Cloning of LCI into YIplac128 was done using Gibson assembly techniques, as described above. The K665 strain was transformed with either the YIplac128 empty vector (EV) or YIplac128 containing LCI using Frozen-EZ Yeast Transformation II Kit (Zymo Research) according to manufacturer protocols and transformed colonies were selected following growth on Synthetic Dropout Leucine (SD-Leu) agar plates overnight at 30°C. Colonies were then grown in SD-Leu liquid media while shaking at 30°C until cultures had an OD of ∼0.6. For the survival assay, a series of 10-fold dilutions of each culture was prepared and dropped onto SD-Leu agar plates with the indicated amounts of added CaCl_2_. After 48h of growth, plates were removed and visualized for growth.

#### Localization in yeast

Cloning of LCI-DsRed into YIplac128 was done using Gibson assembly techniques, as described above. The K665 strain was transformed with either the YIplac128 empty vector (EV) or YIplac128-LCI-DsRed as described above, and transformed colonies were selected following growth on SD-Leu agar plates overnight at 30°C. Colonies were then grown in SD-Leu liquid media while shaking at 30°C until cultures had an OD of ∼0.6. Vacuoles were then labeled with the dye FM4-64, according to manufacturer protocols. Briefly, yeast cultures were harvested by spinning down for 1 minute at 5,000 x g for 5 minutes. The supernatant was discarded and the cell pellet was resuspended in 50 µL Yeast Extract-Peptone-Dextrose (YPD) media and 1 µL of 1.6 µM FM4-64. This mixture was incubated for 20 minutes in a 30°C water bath. Then, 1 mL YPD media was added and the mixture was centrifuged for 5 minutes at 5,000 x g. The supernatant was discarded and the cells were resuspended in 5 mL YPD media. This mixture was incubated in a shaker for 90 minutes at 30°C. The cells were centrifuged for 5 minutes at 5,000 x g before removing the supernatant and resuspending cells in 1 mL sterile water. Cells were then spun down again for 5 minutes at 5,000 x g before resuspending in 25 µL YPD media. On a coated glass slide, 7 µL of the sample was spotted and covered with a coverslip. Cells were then imaged on a widefield microscope.

#### Colocalization in Live Cells

To evaluate the subcellular localization of LCI mutants, we analyzed colocalization with TMR-dextran or FITC-dextran in live cells. In both experiments, COS7 cells were transfected with the indicated variant of LCI-EGFP or LCI-DsRed with Lipofectamine 3000 according to manufacturer protocol and either imaged or fixed 48h later. In the live cell experiments, transfected COS7 cells were pulsed with 1 mg/mL TMR-dextran or 5 mg/mL FITC-dextran for 1h and chased for 16h prior to imaging. Where indicated, COS7 cells were also incubated in 5µM vacuolin-1 for 24h prior to imaging on a confocal microscope. Pearson’s correlation coefficient (PCC) was calculated using software in ImageJ, both before and after a pixel shift. No threshold PCC was used for colocalization with TMR-dextran or FITC-dextran.

#### Immunofluorescence

In anti-GM130 experiments, COS7 cells were transfected with the indicated variant of LCI-EGFP. Two days later, they were washed 3x with PBS and fixed using 4% paraformaldehyde at RT for 10min. Then, cells were washed 3x with PBS and permeabilized with 0.2% Triton X-100 for 5min prior to washing again and blocking with 3% BSA for 60min. Cells were then incubated with 1:1000 anti-GM130 and 1:1000 anti-TMEM165 in 0.3% BSA overnight at 4°C. Finally, cells were washed 3x with PBS and incubated in 1:1000 goat anti-mouse Alexa647 and 1:1000 goat anti-rabbit Alexa488 in 0.3% BSA for 60min before imaging on a confocal microscope. Above threshold PCC was used for colocalization with GM130.

For immunofluorescence to check LCI expression, HeLa cells or LCI KO HeLa cells were fixed, permeabilized, and blocked as above. Cells were then incubated with 1:1000 anti-TMEM165 in 0.3% BSA in PBS overnight at 4°C. Cells were washed 3x with PBS and incubated in 1:1000 goat anti-rabbit Alexa488 in 0.3% BSA in PBS for 60min before imaging on a confocal microscope. Background-subtracted whole cell intensity was used to evaluate expression levels.

For immunofluorescence in cells with swollen lysosomes, HeLa cells were pre-treated with 5µm vacuolin-1 prior to fixation. Two days later, cells were washed 3x with PBS and fixed using 1% glyoxal solution in PBS for 5 minutes. Then, cells were washed 3x with PBS and permeabilized with 0.2% Triton X-100 for 10 minutes. Then, cells were washed 3x with PBS and blocked with 3% BSA for 1 hour. Cells were then incubated in 1:100 mouse anti-LAMP1 and/or 1:200 rabbit anti-TMEM165 in 0,3% BSA overnight at 4°C. Finally, cells were washed 3x with PBS and incubated in 1:1000 goat anti-rabbit Alexa488 and/or 1:1000 goat anti-mouse Alexa647 in 0.3% BSA for 1 hour before imaging on a confocal microscope.

For surface-only immunostaining, HeLa cells were first transfected with LCI-EGFP. Two days later, cells were fixed with 2% PFA for 10 minutes on ice and washed 3x with ice-cold PBS. Cells were then blocked with 3% BSA with 0.3M glycine in PBS for 30 minutes. Cells were then incubated in 1:100 anti-GFP Alexa555 and 1:500 rabbit anti-pan cadherin in blocking buffer overnight at 4°C. After washing 3x with PBS, cells were incubated in 1:1000 goat anti-rabbit Alexa647 in blocking buffer for 1 hour before washing and imaging on a confocal microscope.

#### Lysosomal pH and Ca^2+^ imaging in worms

*In vivo* lysosomal pH and Ca^2+^ measurements were made using *CalipHluor2.0* according to protocols and calibrations established previously^17, 49^. The methods are summarized below.

##### CalipHluor 2.0 preparation

In *CalipHluor 2.0*, the Ca^2+^ indicator, Rhod-5F (O), fluoresces upon binding Ca^2+^[17]. The K_d_ of Rhod-5F for Ca^2+^ binding is pH dependent, which is why *CalipHluor 2.0* incorporates a pH sensing dye dichlorofluorescein (DCF, G)^49^. For ratiometric quantification of Ca^2+^ and pH, we incorporate Alexa Fluor 647 (R) as a reference dye. Thus, the G/R ratio maps the pH at every pixel and is used to obtain pH-corrected values of Ca^2+^ based on the O/R ratio at single-lysosome resolution. *CalipHluor 2.0* was prepared according to previously reported procedure ^49^. Oligonucleotides used to form *CalipHluor 2.0* are listed in the Key Resources Table.

##### In vitro pH and calcium calibration

The calibration curve used here to make pH measurements with the DCF/Alexa647 (G/R) ratio from *CalipHluor 2.0* was prepared in ref. 38. The resulting curve was fitted to Equation 1:

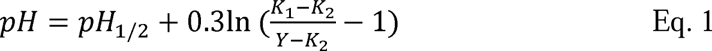

where K_1_, K_2_, and pH_1/2_ represent parameters from a Boltzmann fit of the calibration curve and Y represents the G/R ratio. An *in vitro* bead calibration of *CalipHluor 2.0* at pH 5 on the same day of measurements (see below) was used to correct for day-to-day variation in the calibration curve. The K_d_ curve used to correct for the effect of pH on Rhod-5F binding to Ca^2+^ was determined in ref. 11. The curve was fitted to Equation 2:

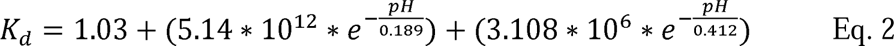

To calculate the effect of pH on the fold-change in the Rhod-5F/Alexa647 (O/R) ratio of *CalipHluor 2.0* from 0.1 µM to 1 mM free Ca^2+^, we used the fold-change calibration curve prepared in ref. 11. The curve was fitted to Equation 3 to get the minimum O/R (O/R at 0.1 µM Ca^2+^) as a function of pH and normalized to the maximum O/R (O/R at 1 mM Ca^2+^):

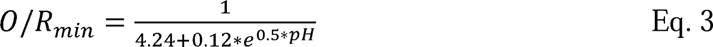

An *in vitro* bead calibration of *CalipHluor 2.0* was performed on the same day as measurements were made to correct for day-to-day variation in fold-change, as previously described ^17, 49^. Briefly, 500 nM of *CalipHluor 2.0* was incubated with 0.6 um monodisperse silica beads in 20 mM sodium phosphate buffer, pH 5.1 containing 500 mM NaCl for 30mins at RT. The beads were washed 3x by spinning at 10k rpm for 10mins at RT. Beads absorbed with *CalipHluor 2.0* were incubated with clamping buffer (HEPES (10 mM), MES (10 mM), sodium acetate (10 mM), EGTA (10 mM), KCl (140 mM), NaCl (500 mM), and MgCl_2_ (1 mM)) for 30 mins at RT containing 0.1 µM or 1 mM free calcium buffers at pH 5. Beads absorbed with *CalipHluor 2.0* were then imaged in DCF (G), Rhod-5F (O), and Alexa647 (R) on a glass slide under the same exposure settings as used for measurements later. Background-subtracted G, O, and R intensities were used to calculate G/R at pH 5.1 to correct Eq. 1 and O/R_min_ and O/R_max_ to correct Eq. 3.

##### In vivo pH and calcium measurements

pH and Ca^2+^ measurements in worms were carried out as previously described for *CalipHluor_Ly_*, but with *CalipHluor 2.0*. *CalipHluor 2.0* is endocytosed by scavenger receptors and marks lysosomes of coelomocytes in live worms^17, 50–52^. Briefly, 500 nM *CalipHluor 2.0* was microinjected into the pseudocoelom of young adult worms with the indicated genetic background. After microinjections, worms were incubated for 2h for maximum labeling of coelomocyte lysosomes. Worms were then anesthetized using 40 mM sodium azide in M9 solution and imaged by wide-field microcopy (details in image acquisition section). Resulting images were background subtracted before calculating the G/R and O/R ratios for each lysosome. Equation 1 was then used to calculate the pH at every lysosome. Equation 2 was then used to calculate the K_d_ of *CalipHluor 2.0* for Ca^2+^ at every lysosome. Equation 3 was then used to calculate the O/R_min_ and O/R_max_ of *CalipHluor 2.0* at every lysosome. Finally, the pH-corrected free [Ca^2+^] was calculated for every lysosome using Equation 4:

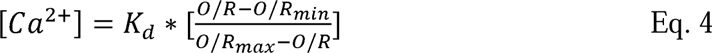

Three independent experiments, each with >5 worms, were made for pH and [Ca^2+^] values for each genetic condition. Lysosomes with O/R values below O/R_min_ were designated as having a [Ca^2+^] less than 0.1 µM. Lysosomes with O/R values above O/Rmax were designated as having a [Ca^2+^] above 1 mM. To estimate a maximum [Ca^2+^] for the exchanger-dead mutant worms, lysosomes with <0.1 µM [Ca^2+^] were counted as having 0.1uM [Ca^2+^].

##### pH and −log[Ca^2+^] maps

Images were acquired in three channels (DCF, Rhod-5F, and Alexa647) by widefield microscopy to quantify pH and [Ca^2+^] at single-lysosome resolution as described above. All image calculations below were done using ImageJ modules. To show representative pH and [Ca^2+^] maps, images were background-subtracted and smoothed. The Alexa647 image was then duplicated and thresholded to create a binary mask. Background-subtracted DCF, Rhod-5F, and Alexa647 images were then multiplied with the binary mask to get processed images. The processed DCF and Rhod-5F images were divided by the processed Alexa647 image to get a pseudocolor G/R and O/R image, respectively. The G/R image was then plugged into Equation 1 to get a pseudocolored pH map. The pseudocolored pH map was then used to get a pseudocolored K_d_ map using Equation 2 and a pseudocolored O/R_min_ map using Equation 3. The O/R image, K_d_ map, and O/R_min_ map were then used to get a [Ca^2+^] map using Equation 4. Finally, to compare maps on an appropriate calibration scale, the −log[Ca^2+^] map was calculated.

#### Lysosomal pH and Ca^2+^ steady-state imaging in cells

Lysosomal pH and Ca^2+^ measurements in live mammalian cells were made using similar methods as in live worms, but with *CalipHluor^mLy^*, which contains pH sensor Oregon Green 488 (OG488) instead of DCF. As such, the preparation and calibration of *CalipHluor^mLy^* were done using the sequences in the Key Resources Table, but with the appropriate calibration curve for OG488, as previously described^17^.

To make measurements in HeLa cells, the appropriate chase time was first determined by analyzing colocalization with endosomal markers. For colocalization with late endosomes, HeLa cells were transfected with Rab7-EGFP. Two days later, cells were treated with 500 nM Alexa647-labeled dsDNA in HBSS for 15 minutes before washing the dsDNA off with PBS and chasing in DMEM for the indicated amount of time (3 hr or 5 hr). For colocalization with lysosomes, HeLa cells were treated with 5 mg/mL FITC-dextran in DMEM for 3 hours. Cells were then washed 3x with PBS and incubated overnight in DMEM. Cells were then treated with 500 nM Alexa647-labeled dsDNA in HBSS for 15 minutes before washing the dsDNA off with PBS and chasing in DMEM for the indicated amount of time (3 hr or 5 hr). For both colocalization experiments, cells were imaged on a confocal microscope in the relevant channels before determining colocalization using Pearson’s Correlation Coefficient (PCC).

pH and Ca^2+^ measurements were then made by incubating HeLa cells in 500 nM *CalipHluor^mLy^* in HBSS for 15 minutes, washing 3x with PBS, and then incubating cells in HBSS for 5 hr. Cells were then imaged on a widefield microscope in the relevant channels. Resulting images were background subtracted before calculating the G/R and O/R ratios for each lysosome. The above equations were used to calculate the pH and Ca^2+^ concentration for individual lysosomes. Three independent experiments, each with >50 lysosomes each, were made for pH and [Ca^2+^] values for each genetic condition. Lysosomes with O/R values below O/R_min_ were designated as having a [Ca^2+^] less than 0.1 µM. Lysosomes with O/R values above O/Rmax were designated as having a [Ca^2+^] above 1 mM. Lysosomes with pH<4 were not included in Ca^2+^ measurements. pH and −log[Ca^2+^] maps were made as described above.

#### Bioinformatics

##### Topology modelling

The secondary structure of LCI was determined using JPred. 2D topology schemes of LCI and vcx1 were then prepared using TOPO2 software (outside residues, membrane residues, signature regions, and special residues off). Transmembrane segments were arranged to display the internal homology of LCI and vcx1 (Tables S2 and S3), with putative Ca^2+^-binding domains forming the pore in the center. Residues are colored to show homology of three critical regions between LCI and vcx-1: Ca^2+^ binding region #1, yellow, 36.36% identity, 54.54% similarity; Ca^2+^ binding region #2, purple, 22.22% identity, 88.88% similarity; acidic helix, green, 31.8% identity, 50.0% similarity. Percentage identity and similarities were calculated using LALIGN (BLOSUM50 matrix; −12 gap open; −2 gap extend; 10.0E() threshold). Light blue indicates the EXGDK/R motifs in LCI and the GNXXE motifs in vcx1, which are involved in Ca^2+^ binding. Red indicate the lysosome-targeting YNRL motif in LCI and the vacuole-targeting YNRV motif in vcx.1

##### Sequence alignments

Alignments of vcx1, TMEM165, gdt1, and Y54F10AL.1 sequences were performed using Clustal Omega (output format: ClustalW with character counts; mBed-like clustering; 0 combined iterations; guide tree iterations off; HMM iterations off). Select regions were highlighted based on homology identified above and outlined in the respective colors.

##### Homology modelling

Homology-based modelling of LCI was performed using MODELLER using vcx1 as a unifying template; information about other templates is provided in Table S4. The script was obtained from the MODELLER online manual provided by A. Sali using the automodel class. The resulting model was uploaded into PyMol and colored according to the previous designations.

#### Cytosolic Ca^2+^ and lysosomal pH/Ca^2+^ dynamics

To evaluate cytosolic Ca^2+^ dynamics, we used the ratiometric Ca^2+^ dye Fura Red according to manufacturer protocols. HeLa cells were pulsed with 10 µM Fura Red in HBSS for 15 min, washed 3x with PBS, and then chased for 30min in HBSS. Prior to imaging in Tyrode’s solution (NaCl 134 mM, KCl 2.68 mM, CaCl_2_ 1.8 mM, MgCl_2_ 1.05 mM, NaH_2_PO_4_ 0.417 mM, NaHCO_3_

11.9 mM, d-glucose 5.56 mM), cells were washed 3x again with PBS. The imaging protocol was set up to Fura Red images every 5 seconds. After 8 image acquisitions, the solution was replaced with Tyrode’s solution supplemented with 100 µM ATP. Images were taken for 5 minutes. Fura Red images were then background-subtracted by the intensity of an ROI outside the cell. The 440/647 image was then duplicated and thresholded to create a binary mask. Both images were multiplied by the mask to create processed 440/647 and 488/647 images. The processed 440/647 image was then divided by the processed 488/647 image to get a 440/488 (excitation ratio) pseudocolored map. For representative images, the pseudocolored maps at indicated time points were then smoothed. The 440/488 ratio was then plotted as a function of time, normalized to 1 at t=0s.

To evaluate lysosomal pH population dynamics, we used the ratiometric lysosomal pH indicator FITC-dextran according to manufacturing protocols. HeLa cells were pulsed with 5 mg/mL FITC-dextran for 3 hours in DMEM before washing 3x with PBS and incubating for 16 hours in DMEM. Prior to imaging in Tyrode’s solution, cells were washed 3x again with PBS. The imaging protocol was set up to take FITC-dextran images every 5 seconds. After 8 image acquisitions, the solution was replaced with Tyrode’s solution supplemented with 100 µM ATP. Images were taken for 5 minutes. FITC-dextran images were then background-subtracted by the intensity of an ROI outside the cell. The 440/514 image was then duplicated and thresholded to create a binary mask. Both images were multiplied by the mask to create processed 440/514 and 488/514 images. The processed 488/514 image was then divided by the processed 440/514 image to get a 488/440 (excitation ratio) pseudocolored map. For representative images, the pseudocolored maps at indicated time points were then smoothed. The 488/440 ratio was then plotted as a function of time, normalized to 1 at t=0s.

To evaluate the effect of LCI on single-lysosome pH and Ca^2+^ dynamics, we used *CalipHluor^mLy^* to generate two-ion measurement maps before and after a cytosolic Ca^2+^ spike. To label lysosomes, WT or LCI KO HeLa cells were pulsed with 500 nM *CalipHluor^mLy^*in HBSS for 15 minutes, washed 3x with PBS, and incubated cells in HBSS for 5 hr. Cells were then imaged on a widefield microscope (see image acquisition section) in the relevant channels before incubating cells in HBSS with 100 µM ATP. Two minutes later, cells were imaged again in the same channels. Resulting pre-ATP and post-ATP images were background subtracted before calculating the G/R and O/R ratios for each lysosome. The above equations were used to calculate the pH and Ca^2+^ concentration for individual lysosomes before and after ATP addition. Lysosomes with O/R values above O/Rmax were designated as having a [Ca^2+^] above 1 mM. Lysosomes with pH<4 were labeled as such, and not included in Ca^2+^ measurements. pH and −log[Ca^2+^] maps were made as described above. A lysosomal pH increase or decrease was defined as an increase or decrease, respectively, of 0.1 pH units or more. A lysosomal Ca^2+^ increase or decrease was defined as a fold change of more than 1.5 or less than 0.67, respectively.

#### Whole-cell electrophysiology

Whole-cell recordings were performed with an Axopatch 200 A amplifier (Molecular Devices) and digitized using an NI-6251 DAQ (National Instruments). The amplifier and digitizer were controlled using WinWCP software (Strathclyde Electrophysiology Software). All data were sampled at 10 kHz and later filtered at 1 kHz using a 4-pole lowpass Bessel filter. The borosilicate glass capillaries (Sutter) with dimensions of 1.5 mm x 0.86 mm (OD/ID) were pulled using a Sutter P-97 micropipette puller and polished using Coater/polisher Microforge (ALA Scientific Instruments Inc.) to achieve a resistance of 2-4 MΩ. Patch pipettes were then positioned using an MP325 motorized manipulator (Sutter). The buffers used for the bath and pipette solutions were designed to reduce background currents from ions other than calcium. The extracellular/bath solution contained (in mM): 140 NMDG, 10 HEPES, 1 MgCl_2_, 5 EGTA, 5 glucose, and variable Ca(OH)_2_ to get 30 nM, 300 nM, or 100 µM free [Ca^2+^], and was set to either pH 5.5, 6.5, or 7.5 with MSA. The pipette solution contained (in mM): 140 NMDG, 10 HEPES, 1 MgCl_2_, 5 EGTA, and variable Ca(OH_2_) to get either 30 nM, 100 nM, 1 µM, or 300 nM free [Ca^2+^], and was set to pH 5.5 or 7.5 with MSA. Total calcium at each pH was calculated to maintain the indicated amount of free calcium using https://somapp.ucdmc.ucdavis.edu/pharmacology/bers/maxchelator/CaMgATPEGTA-TS.htm for each experiment. Where indicated, internal Ca(OH)_2_ was replaced with Ba(OH)_2_ or MgCl_2_ to get the equivalent concentration of divalent cation. Mock-transfected or LCI-EGFP (WT or E248A)-transfected HeLa cells were washed in PBS before incubating in the indicated extracellular solution for whole-cell clamping. Once the whole-cell configuration was established, membrane capacitance and series resistance were compensated according to established protocols ^53^. A ramp protocol (−100mV to +100mV, 1sec, holding at 0mV) was used to record the currents. All experiments were performed at room temperature (22-23°C) and analyzed with WinWCP and OriginPro 2022b (Origin Lab). As indicated in Figure S6B, positive voltage is taken as cytosol-positive, and positive current is taken as positive current moving outward from the cytosol. Current density was calculated by normalizing the current with whole-cell capacitance, which was determined immediately after establishing the whole-cell patch to estimate cell surface area. Average traces shown were smoothed using 7-point adjacent-averaging. Where indicated, the current density at +100mV or −100mV was plotted for each condition. Theoretical reversal potentials were calculated as previously described^54^.

#### Lysosome electrophysiology

Borosilicate glass capillaries (Sutter) with dimensions of 1.5 mm x 0.86 mm (outer diameter/inner diameter (OD/ID)) were pulled using the following program: heat, 520; pull, 0; vel, 20; time, 200; loops, 4. Pipettes were fire-polished using an MF200 microforge (World Precision Instruments). Fire-polished patch pipettes were then used in voltage-clamping experiments. For the lysosome patching experiment shown in Figure S7D-E, the pipette and bath solutions were designed to mimic the ionic composition of the lysosome (pH 4.5) or Golgi (pH 6.2) and cytoplasm, respectively, and are shown in Figure S7C. The cytosolic/batch solution contained (in mM): 20 KCl, 120 K-gluconate, 2 MgCl_2_, 2.5 CaCl_2_, 0.2 EGTA, 10 HEPES, 2 Na_2_ATP, pH 7.25. The pipette/lumenal solution contained (in mM): 145 NaCl, 20 glucose, 5 Na3Cit, 10 EGTA, 2 CaCl2, 1 MgCl2, 10 TEA, 3 KCl, pH 4.5 or 6.2. Mock-transfected or LCI-EGFP-transfected COS7 cells were treated with 5 µM vacuolin-1 overnight to increase the side of lysosomes to 1-3 µm ^55^. Cells were then washed 3x with PBS before being incubated in the indicated bath solution. Enlarged lysosomes were pushed out of the ruptured cell. After giga-ohm seal formation, break-in was performed by a zap protocol (5 V: 0.5-5s) until the appearance of capacitance transients. Voltage ramping and calculation of current were performed as described above. As indicated in Figure S7C, positive membrane potential is taken as cytosol-positive, and positive current is taken as positive current moving outward from the cytosol. Representative traces shown were smoothed using 24-point adjacent-averaging. Where indicated, the current at +100 mV was plotted for each condition.

#### Image acquisition

Widefield microscopy was carried out on an IX83 inverted microscope (Olympus Corporation of the Americas, Center Valley, PA, USA) using a 60X, 1.4 numerical aperture (NA), differential interference contract (DIC) oil immersion objective (PLAPON) and Evolve Delta 512 EMCCD camera (Photometrics, USA), and controlled using MetaMorph Premier Ver 7.8.12.0 (Molecular Devices, LLC, USA). For *CalipHluor2.0* imaging in worms, images were acquired with an exposure of 150ms and EM gain of 150 for DCF, an exposure of 150ms and EM gain of 150 for Rhod-5F, and an exposure of 50ms and EM gain of 50 for Alexa647. DCF channel images were obtained using a 480/20 band-pass excitation filter, 520/40 band-pass emission filter, and an 89016 dichroic; Rhod-5F channel images were obtained using a 545/25 band-pass excitation filter, 595/50 band-pass emission filter, and an 89016 dichroic; and Alexa647 channel images were obtained using a 640/30 band-pass excitation filter, 705/72 band-pass emission filter, and an 89016 dichroic. For *CalipHluor^mLy^* imaging in cells, images were acquired with an exposure 200ms and EM gain of 200 for OG488, an exposure of 200ms and EM gain of 200 for Rhod-5F, and an exposure of 200ms and EM gain of 200 for Alexa647. OG488 channel images were obtained using a 480/20 band-pass excitation filter, 520/40 band-pass emission filter, and an 89016 dichroic; Rhod-5F channel images were obtained using a 545/25 band-pass excitation filter, 595/50 band-pass emission filter, and an 89016 dichroic; and Alexa647 channel images were obtained using a 640/30 band-pass excitation filter, 705/72 band-pass emission filter, and an 89016 dichroic. For cytosolic Ca^2+^ recording cells, Fura Red was recorded by excitation at 440nm or 488nm and emission at 647nm. 440/647 images were acquired using a 430/24 band-pass excitation filter, 705/72 band-pass emission filter, and an 89016 dichroic with an exposure of 150ms and EM gain of 150. 488/647 images were acquired using a 480/20 band-pass excitation filter, 705/72 band-pass emission filter, and an 89016 dichroic with an exposure of 200ms and EM gain of 200. For lysosomal pH recording in cells, FITC-dextran was recorded by excitation at 440nm or 488nm and emission at 514nm. 440/514 images were acquired using a 430/24 band-pass excitation filter, 520/40 band-pass emission filter, and an 89007 dichroic with an exposure of 200ms and EM gain of 200. 488/514 images were acquired using a 480/20 band-pass excitation filter, 520/40 band-pass emission filter, and an 89016 dichroic with an exposure of 200ms and EM gain of 200. EGFP images were acquired using a 480/20 band-pass excitation filter, 520/40 band-pass emission filter, and an 89016 dichroic. DsRed images were acquired using a 545/25 band-pass excitation filter, 595/50 band-pass emission filter, and an 89016 dichroic. FM4-64 images were acquired using a 545/25 band-pass excitation filter, 632/60 band-pass emission filter, and an 89016 dichroic.

Confocal images were captured with a Leica TCS SP5 II STED laser confocal microscope (Leica Microsystems, Buffalo Grove, IL, USA) equipped with a 63X, 1.4 NA, oil immersion objective. GFP and Alexa488 were excited using an argon laser with wavelength of 488nm; Alexa647 was excited using an He-Ne laser with wavelength of 633nm; and TMR, DsRed, and Alexa555 were excited using a DPSS laser at 561nm. All emissions were filtered using an acousto-optical beam splitter (AOBS) with settings suitable for each fluorophore and recorded using hybrid detector.

#### Image analysis

Images were analyzed using Fiji (NIH, USA). For lysosomal pH and Ca^2+^ measurements with *CalipHluor 2.0*, regions of coelomocytes containing single isolated lysosomes in each Alexa647 (R) image were manually selected and the coordinates were saved in the ROI plugin. For each coelomocyte, the most focused plane was manually selected in the Alexa647 channel and used for all other channels. For background computation, a nearby region outside of the worm was manually selected and saved as an ROI. The same regions were selected in the DCF (G) and Rhod-5F (O) images by recalling the ROIs. After background subtraction, the mean intensity for each endosome (G, O, and R) was measured and exported to OriginPro (OriginLab, USA). Ratios of G to R intensities (G/R) and O to R intensities (O/R) were obtained by dividing the mean intensity of a given lysosome in the G or O image by the corresponding intensity in the R image. Representative images are shown as pseudo-colored maps, where G, O, and R images were modified by thresholding in ImageJ before dividing and undergoing the relevant image calculations in ImageJ. Analogous image analysis methods were used to make lysosomal Ca^2+^ measurements in cells with *CalipHluor^mLy^*, lysosomal pH measurements in cells with FITC-dextran and cytosolic Ca^2+^ measurements in cells with Fura Red.

### Quantification and Statistical Analysis

All data are presented with statistics indicated in the relevant figure legend. For statistical analysis between two samples, a two-sample two-tailed test assuming unequal variance was conducted. For comparison of multiple samples, one-way ANOVA with a post hoc Tukey test was conducted. All statistical analysis was performed in Origin Lab.

## Supplementary Information Titles and Legends

**Table S1**

Genes screened for *cup-5^+/-^*worm survival rescue, with brood size difference and significance calculated with respect to empty vector (EV).

