## Supplementary material for "TMEM165 acts as a proton-activated Ca^2+^ importer in lysosomes": Key Resources Table

| REAGENT or RESOURCE | SOURCE | IDENTIFIER |
| --- | --- | --- |
| Antibodies | | |
| Rabbit anti-TMEM165 | Proteintech | Cat #: 20485-1-AP |
| Mouse anti-GM130 | Santa Cruz Biotechnology | Cat #: SC-55591 |
| Alexa488 goat anti-rabbit | Thermo Fisher Scientific | Cat#: A1101 |
| Alexa 647 goat anti-mouse | Thermo Fisher Scientific | Cat#: A21244 |
| GFP polyclonal antibody Alexa555 | Thermo Fisher Scientific | Cat#: A-31851 |
| Anti-pan cadherin | Abcam | Cat#: ab16505 |
| Bacterial and virus strains | | |
| DH5α E. coli | NEB | N/A |
| OP50 | CGC | N/A |
| Chemicals, peptides, and recombinant proteins | | |
| EGTA | Sigma | Cat #: 324626 |
| Ampicillin | Fisher | Cat #: BP1760 |
| Kanamycin | Sigma | Cat #: 60615 |
| Carbenicillin | Sigma | Cat #: C9231 |
| Adenosine triphosphate | Sigma | Cat #: A26209 |
| Calcium chloride | Sigma | Cat #: 223506 |
| SD-Trp | Sigma | Cat #: Y1876 |
| SD-Leu | Sigma | Cat #: Y1376 |
| FITC-dextran (10 kDa) | Sigma | Cat #: FD10S |
| IPTG | Sigma | Cat #: I6758 |
| YNB with ammonium sulfate | MPBio | Cat #: MP114027512 |
| TMR-dextran (10 kDa) | Thermo Fisher Scientific | Cat #: D1816 |
| Fura Red | Thermo Fisher Scientific | Cat #: F3021 |
| DMEM | Life Technologies | Cat #: 12400024 |
| FBS | Gibco | Cat #: 26140079 |
| Penstrep | Life Technologies | Cat #: 15140122 |
| Trypsin-EDTA | Life Technologies | Cat #: 25200056 |
| LongAmp Taq DNA Polymerase | NEB | Cat #: M0323S |
| DpnI | NEB | Cat #: R0176S |
| DNA Clean and Concentrator Kit | Zymo Research | Cat #: D4003 |
| Gibson Assembly Master Mix | NEB | Cat #: E2611S |
| Luria Broth | Fisher | Cat #: BP9723 |
| GeneJET Plasmid Miniprep Kit | Thermo Fisher Scientific | Cat #: K0502 |
| Q5 Site-Directed Mutagenesis Kit | NEB | Cat #: E0554S |
| Lipofectamine 3000 | Thermo Fisher Scientific | Cat #: L3000-001 |
| Frozen-EZ Yeast Transformation II Kit | Zymo Research | Cat #: T2001 |
| Agarose | Gold biotech | Cat #: A-201-500 |
| Tris base | Fisher Scientific | Cat #: BP152 |
| EDTA | Sigma | Cat #: E5134 |
| PFA | Fisher Scientific | Cat #: T353 |
| Triton X-100 | Sigma | Cat #: T8787 |
| BSA | Sigma | Cat #: A7906 |
| HEPES | Sigma | Cat #: H3375 |
| MES | Sigma | Cat #: M3671 |
| Sodium acetate | Fisher Scientific | Cat #: S210 |
| Potassium chloride | Fisher Scientific | Cat #: BP366 |
| Sodium chloride | Alfa Aesar | Cat #: A12313 |
| Magnesium chloride | Sigma | Cat #: M8266 |
| Sodium phosphate monobasic | Sigma | Cat #: S3139 |
| Sodium bicarbonate | Fisher Scientific | Cat #: S233 |
| Dextrose | Fisher Scientific | Cat #: D16 |
| NMDG-Cl | TCI | Cat #: M0713 |
| TEA-Cl | TCI | Cat #: T0095 |
| Sodium methanesulfonate | Acros | Cat #: 442110050 |
| HBSS | Fisher Scientific | Cat #: 14025092 |
| Trizol | Life Technologies | Cat #: 15596026 |
| Chloroform | Sigma | Cat #: 288306 |
| Isopropanol | Sigma | Cat #: I9516 |
| Superscript IV Reverse Transcriptase | Fisher Scientific | Cat #: 18090010 |
| Vacuolin-1 | Cayman Chemical | Cat#: 20425 |
| Glyoxal | Sigma | Cat#: 128465 |
| Methane sulfonic acid | Sigma | Cat#: 471356 |
| Calcium hydroxide | Sigma | Cat#: 239232 |
| Barium hydroxide | Sigma | Cat#: 433373 |
| Ammonium chloride | Sigma | Cat #: A9434 |
| Experimental models: Cell lines | | |
| HeLa | ATCC | Cat #: CCL-2 |
| COS7 | ATCC | Cat #: CRL-1651 |
| Experimental models: Organisms/strains | | |
| C. elegans wild isolate | CGC | *N2* |
| C. elegans: *+/mT1 II; cup-5(ok1698)/mT1 [dpy-10(e128)] III* | CGC | *VC1242 (cup-5^+/-^)* |
| C. elegans: *arIs37[myo-3p::ssGFP+dpy-20(+)]I* | CGC | *arIs37* |
| C. elegans: *arIs37[myo-3p::ssGFP+dpy-20(+)]Icup5(ar465)* | CGC | *GS2477 (arIs37; cup5(ar465))* |
| C. elegans: *Y54F10AL.1(gk5484[loxP + Pmyo-2::GFP::unc-54 3' UTR + Prps-27::neoR::unc-54 3' UTR + loxP])/+ III* | Moermen Laboratory | *lcax-1*^+/-^ |
| *lcax-1*^+/-^ + Ex: PY54 F10AL.1-LCAX-1-unc-54 3'UTR | Suny Biotech | WT |
| *lcax-1*^+/-^ + Ex: PY54 F10AL.1-LCAX-1 (R126C)-unc-54 3'UTR | Suny Biotech | R126C |
| *lcax-1*^+/-^ + Ex: PY54 F10AL.1-LCAX-1 (G304R)-unc-54 3'UTR | Suny Biotech | G304R |
| *lcax-1*^+/-^ + Ex: PY54 F10AL.1-LCAX-1 (E108A)-unc-54 3'UTR | Suny Biotech | E108A |
| *lcax-1*^+/-^ + Ex: PY54 F10AL.1-LCAX-1 (E188A)-unc-54 3'UTR | Suny Biotech | E188A |
| *lcax-1*^+/-^ + Ex: PY54 F10AL.1-LCAX-1 (E248A)-unc-54 3'UTR | Suny Biotech | E248A |
| S. cerevisiae: Δvcx1Δpmc1 | Hirschi Laboratory | K665 |
| Oligonucleotides | | |
| DBCO-ATCAACACTGCACACCAGACAGCAAGATCCTATATATA | IDT | D1 |
| Alexa 647-TATATATAGGATCTTGCTGTCTGGTGTGCAGTGTTGAT | IDT | D2 |
| Amino-ATAACACATAACACATAACAAAATATATATCCTAGAACGAC AGACAAACAGTGAGTC | IDT | C1 |
| ATTO647-TATATTTTGTTATGTGTTATGTGTTAT | IDT | C2 |
| DBCO-GACTCACTGTTTGTCTGTCGTTCTAGGATA | IDT | C3 |
| Oregon Green- ATAACACATAACACATAACAAAATATATATCCTAGAACGACAGACAAACAGTGAGTC | IDT | OG-C1 |
| GCGCTATAACTGCCTGACCGT | IDT | R126C_F |
| ATTGCCATGATGGCTGCTATAAAAAATG | IDT | R126C_R |
| CGCTATAACCACCTGACCGTG | IDT | R126H_F |
| CATTGCCATGATGGCTGC | IDT | R126H_R |
| GACAATCATAAGAGGCATCGTTTTTTTG | IDT | G304F_F |
| ACAGTTCTGACAGAGATTTTC | IDT | G304F_R |
| L4440 RNAi vectors | Ahringer Library | N/A |
| YIplac128 | Glick Laboratory | N/A |
| pEGFP-N1 | Fransen Laboratory | N/A |
| ATTCCACCGATTTCCACCCC | IDT | Y54F10AL.1_F |
| CTTCTTGGCCCTCATTCGGT | IDT | Y54F10AL.1_R |
| ACAACTACAAATGCGATGACCA | IDT | ncx-1_F |
| ATTGTCGATGGTCCCAGCTC | IDT | ncx-1_R |
| TTCGCTACCATCCCACCAAC | IDT | ncx-2_F |
| CAGTAGGCTTCCAATGCGGA | IDT | ncx-2_R |
| CGGTTTGGTGACTGCTGTTG | IDT | ncx-3_F |
| CGTAGACAATCCAGAGGCCC | IDT | ncx-3_R |
| GTCTACCGATTCCGTGGCTT | IDT | ncx-4_F |
| CCGTTGATGCAGACCGTTTT | IDT | ncx-4_R |
| ACGACTGCAGGGTGTATGTG | IDT | clh-6_F |
| CGAGACCTGTCCATGAGAGC | IDT | clh-6_R |
| ATTGTTGGTGCAGTCAACGC | IDT | catp-6_F |
| GGGGGAATGTTATGCAAGTCG | IDT | catp-6_R |
| GCGGTTAGAGCAAATTCCCC | IDT | cup-5_F |
| TGAGCGCCAAGATTTCCAGA | IDT | cup-5_R |
| CACCAGCAGCTCCAGTTCAT | IDT | TMEM165_F |
| TGAGAGCGATCACCCCATTC | IDT | TMEM165_R |
| TGCAGAAGGAAATCACCGCT | IDT | act-1_F |
| AGAAAGCTGGTGGTGACGAT | IDT | act-1_R |
| Recombinant DNA | | |
| TMEM165 cDNA | Harvard Plasmid Database | HsCD00332210 |
| Software and algorithms | | |
| Fiji | NIH | https://imagej.net/Fiji |
| OriginPro | OriginLab | https://www.originlab.com/ |
| JPred | University of Dundee | http://www.compbio.dundee.ac.uk/jpred/ |
| LALIGN | SIB | https://embnet.vital-it.ch/software/LALIGN_form.html |
| Clustal Omega | EMBL-EBI | https://www.ebi.ac.uk/Tools/msa/clustalo/ |
| WinWCP | University of Strathclyde | http://spider.science.strath.ac.uk/sipbs/software_ses.htm |
| MetaMorph Premier Ver 7.8.12.0 | Molecular Devices | https://www.moleculardevices.com/products/cellular-imaging-systems/acquisition-and-analysis-software/metamorph-microscopy#gref |
| MODELLER | Sali Lab | https://salilab.org/modeller/ |
| Other | | |
| Monodisperse silica microspheres | Cospheric | Cat #: SiO2MS-2.0 0.690um - 1g |
| IX83 inverted microscope | Olympus | https://www.olympus-lifescience.com/en/microscopes/inverted/ix83/ |
| Borosilicate glass capillaries | Sutter | Cat #: BF150-86-10 |
| Leica SP5II confocal microscope | Leica | https://www.leica-microsystems.com/ |
| Axopatch 200 A amplifier | Molecular Devices | https://www.moleculardevices.com/products/axon-patch-clamp-system/amplifiers/axon-instruments-patch-clamp-amplifiers#gref |
| Sutter P-97 Micropipette Puller | Sutter | https://www.sutter.com/MICROPIPETTE/p-97.html |
| MP325 motorized manipulator | Sutter | https://www.sutter.com/MICROMANIPULATION/mpc325_frame.html |
| SZX-Zb12 Stereomicroscope | Olympus | https://www.olympus-ims.com/en/microscope/szx/ |
| MF200 microforge | World Precision Instruments | https://www.wpiinc.com/var-3114-analog-microforge |
| NI-6351 DAQ Digitizer | National Instruments | https://www.ni.com/en-us/shop/data-acquisition.html |
