## Supplementary Information for "TMEM165 acts as a proton-activated Ca^2+^ importer in lysosomes"

**Supplementary Figures**

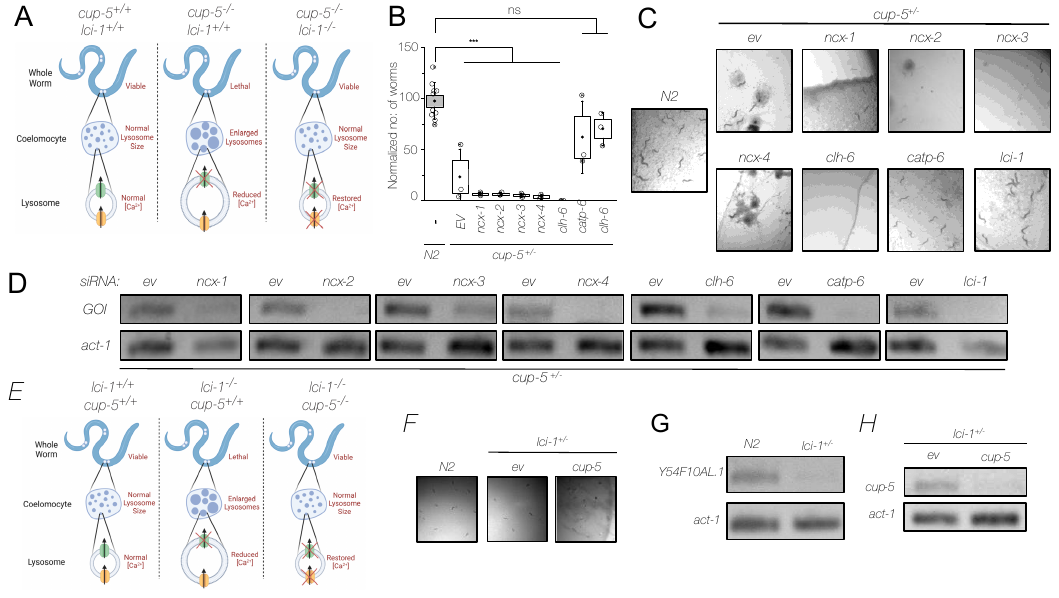

**Figure S1 | The activity of ­*lci-1* complements *cup-5***.

(A) Schematic of the principle underlying screening for lysosomal calcium importers (LCIs) in cup-5^+/-^ worms by survival and lysosome size.

(B) Number of *cup-5*^+/-^ progeny following RNAi knockdown of indicated transcripts.

(C) Representative images showing the number of progeny of N2 worms or *cup-5^+/-^* worms in plates containing RNAi bacteria of empty vector (EV, control), *ncx-1*, *ncx-2*, *ncx-3*, *ncx-4*, *clh-6*, *lci-1*, or *catp-6* (positive control).

(D) RT-PCR analysis of total RNA isolated from *cup-5^+/-^* worms for the indicated gene of interest (GOI) following knockdown of the indicated gene, compared to treatment with empty vector. The activity of ­*cup-5* complements *lci-1.*

(E) Schematic of principle behind phenotypes observed in *lci-1*^+/-^ worms.

(F) Representative images showing the number of progeny of N2 worms or *lci-1*^+/-^ worms in plates containing RNAi bacteria of empty vector (ev, control) or ­*cup-5*.

(G) RT-PCR analysis of total RNA isolated from N2 or *lci-1*^+/-^ worms for the ­*lci-1* (*Y54F10AL.1)* gene, with actin used as a control.

(H) RT-PCR analysis of total RNA isolated from *lci-1*^+/-^ worms for the *cup-5* gene following knockdown of ­*cup-5*, compared to treatment with empty vector.

***p<0.001 (one-way ANOVA with Tukey post hoc test).

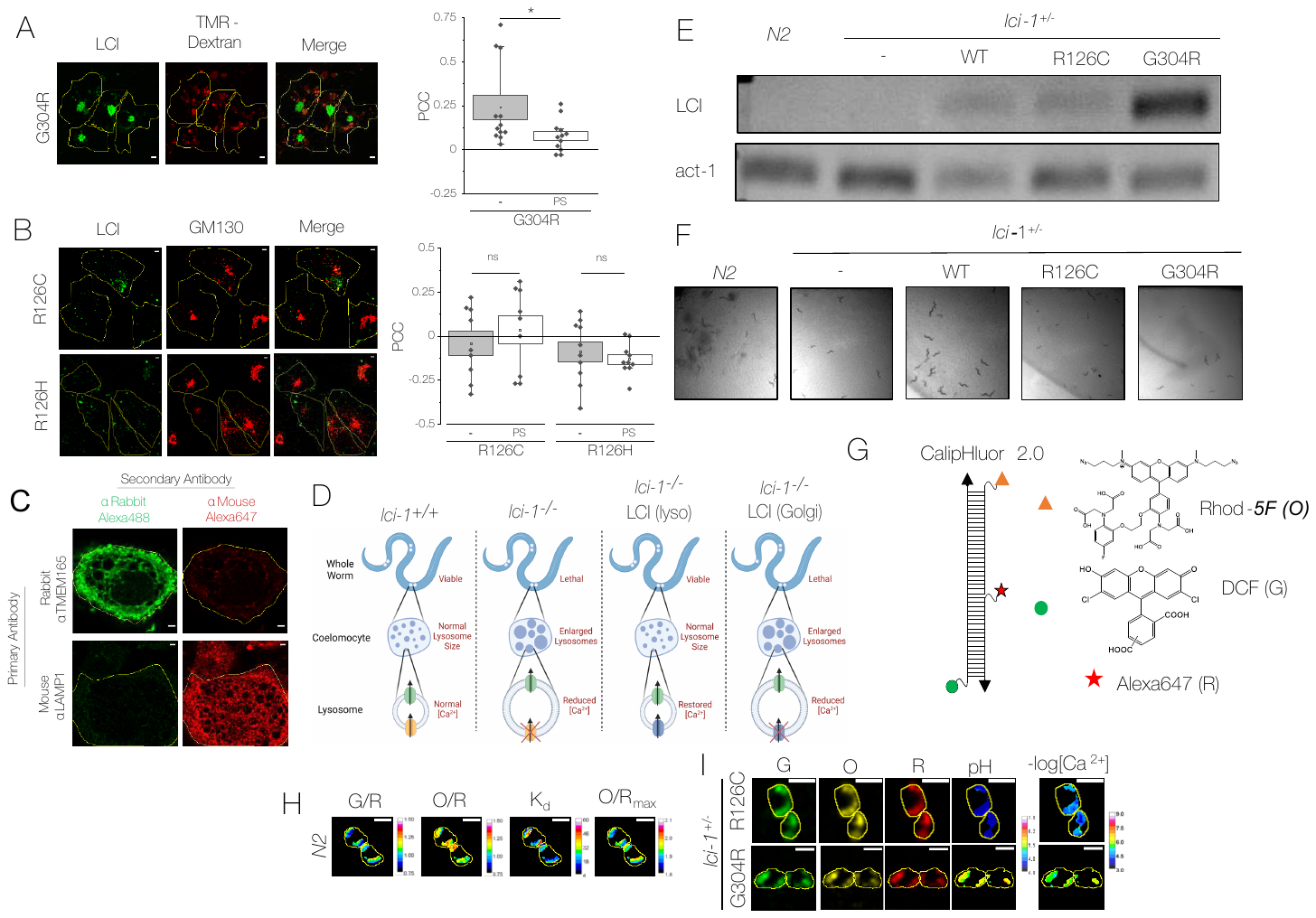
**Figure S2 | Localization of LCI and phenotypic assays in *lci-1*^+/-^ worms.**

(A) Left, representative fluorescence images of COS-7 cells transiently expressing G304R LCI-EGFP (green) and labeled with TMR-dextran (red). Right, Pearson correlation coefficient (PCC) of colocalization between G304R LCI and TMR-dextran before and after pixel shift (PS).

(B) Left, representative fluorescence images of COS-7 cells transfected with the indicated mutant of LCI and immunostained for LCI (green) and GM130 (red). Right, pearson correlation coefficient (PCC) of colocalization between LCI mutants and GM130 before and after pixel shift (PS). Lysosomal localization of LCI.

(C) Representative immunofluorescence images showing specificity of secondary antibodies and spectral separation of fluorophores used in Figure 2E.

(D) Schematic of principle underlying rescue of *lci-1^+/-^* worm phenotypes with human LCI variants.

(E) RT-PCR analysis of the LCI (TMEM165) expression in N2 worms and *lci-1*^+/-^ worms with extrachromosomal expression of the indicated mutant of human LCI.

(F) Representative images showing the number of progeny of N2 worms or *lci-1*^+/-^ worms with extrachromosomal expression of the indicated mutant of human LCI.

(G) Schematic of *CalipHluor 2.0*, which consists of Rhod-5F (orange triangle), DCF (green circle), and Alexa647 (red star) on a DNA duplex.

(H) Representative pseudocolored maps of the DCF/Alexa647 ratio (G/R) and Rhod-5F ratio (O/R) of *CalipHluor2.0* in N2 worms. These maps are used according to equations in the Methods section to generate K_d_ and O/R_max_ maps.

(I) Representative fluorescence images and pH and -log([Ca^2+^]) maps in *CalipHluor2.0*-labeled lysosomes in coelomocytes in *lci-1*^+/-^ worms extrachromosomally expressing the indicated variant of LCI. G, DCF; O, Rhod-5F; R, Alexa647.

Scale bar 5 µm. Inset scale bar 2um. Boxes and bars represent the s.e.m. and standard deviation, respectively. ns, not significant (P>0.05); *P<0.05.

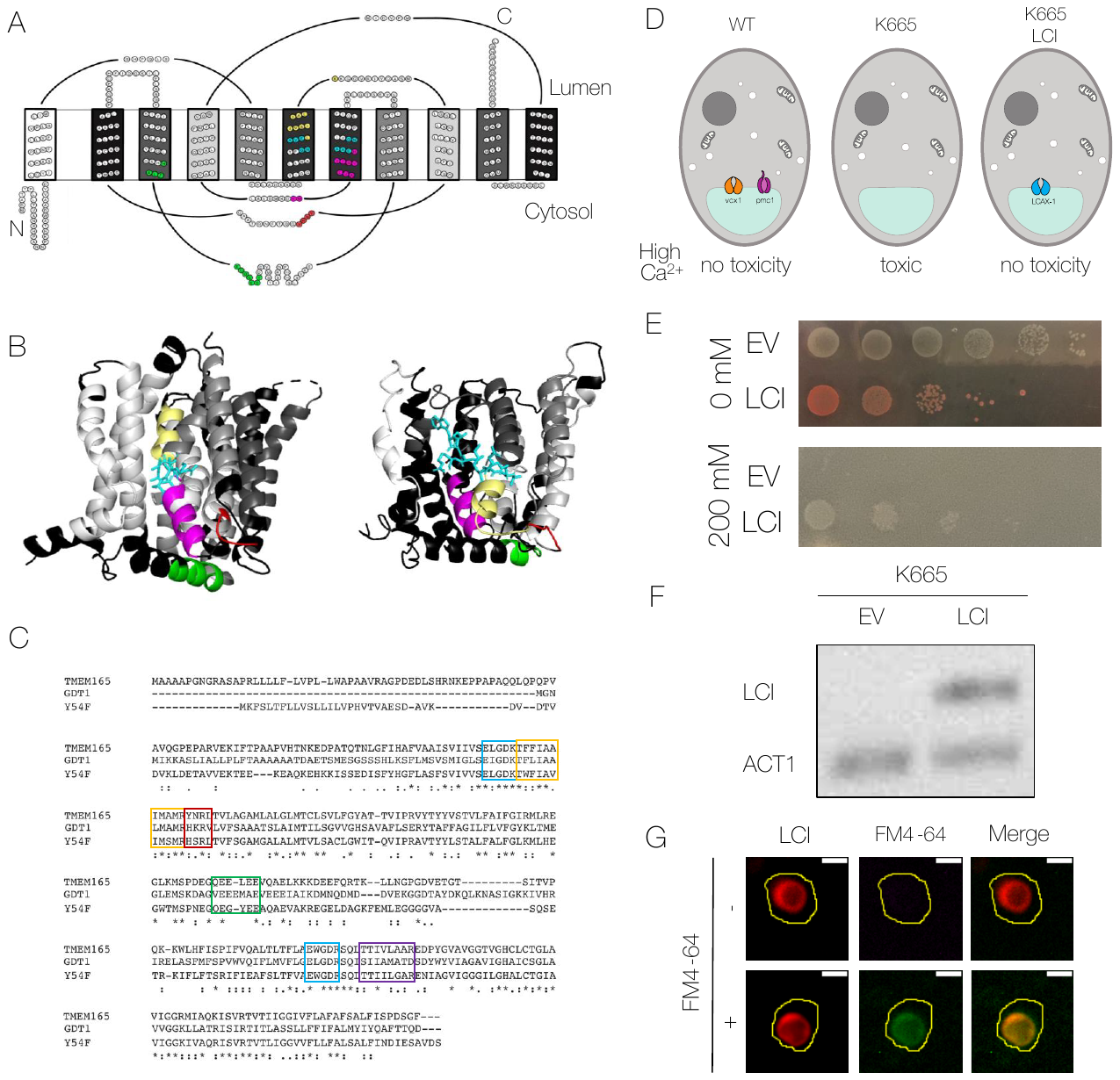
**Figure S3 |** **Sequence and functional conservation within the UPF0016 family**.

(A) Topology of vcx1 created using TOPO2. Grayscale shows internal symmetry of transmembrane segments. Ca^2+^ binding regions (light blue), homologous regions adjoining the Ca^2+^ binding regions (yellow and purple), acidic helices (green), and lysosome/vacuole-targeting motifs (red) are highlighted.

(B) Full sequence alignment of three proteins in the UPF0016 family: TMEM165 (*H. sapiens*), Y54F10AL.1 (*C. elegans*), and gdt1 (*S. cerevisiae*) using Clustal Omega. Asterisks indicate full conservation, colons indicate strong conservation, and periods indicate weak conservation. Calcium binding regions, homologous regions to vcx1, acidic helices, and lysosome/vacuole-targeting motifs are highlighted according to colors indicated in Figure 4.

(C) Homology-based model of LCI (right) based on vcx1 (left), with colors highlighted according to Figure 4 indications.

(D) The rescue assay in *S. cerevisiae* strain K665, where the loss of vcx1 and pmc1 cause lethality in environmental Ca^2+^*.* Expression of a vacuolar Ca^2+^ importer would rescue this lethality at high Ca^2+^ by restoring vacuolar function.

(E) Color of K665 transformed to integrate an empty vector (EV) or human LCI, after 2 days at 30°C on YPD plates supplemented with the indicated concentration of CaCl_2_. Columns indicated 10-fold dilutions from left-to-right.

(F) RT-PCR of TMEM165 (*H. sapiens*) and ACT1 (*S. cerevisiae*) in the K665 strain transformed with empty vector (EV) or LCI.

(G) Representative fluorescence images of K665 transformed with LCI-DsRed (red) and labelled with the vacuolar membrane marker FM4-64 (green), where indicated.

Scale bar 2µm.

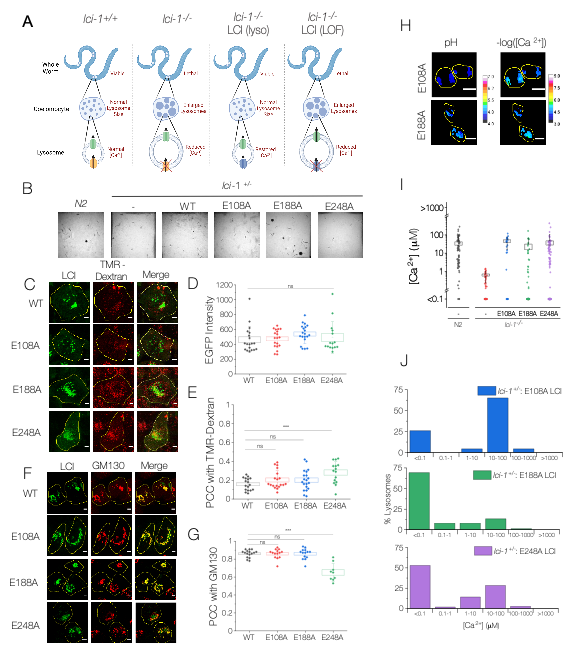

**Figure S4 | Functional LCI mutants rescuing organism and cellular phenotypes seen in *lci-1^+/-^* worms.**

(A) Schematic of principle underlying rescue of *lci-1^+/-^* worm phenotypes with human LCI loss-of-function (LOF) variants.

(B) Representative images showing the number of progeny of N2 worms or *lci-1*^+/-^ worms with extrachromosomal expression of the indicated mutant of human LCI.

(C) Representative fluorescence images of HeLa cells transfected with the indicated LCI mutant (green) and labeled with 5 mg/mL TMR-dextran (red).

(D) Whole-cell fluorescence intensity of LCI-EGFP in cells transfected with the indicated mutant. (E) Pearson correlation coefficient (PCC) of LCI-EGFP with TMR-dextran in cells transfected with the indicated mutant.

(F) Representative immunofluorescence images of HeLa cells transfected with the indicated LCI mutant (green) and stained with an antibody to GM-130 (red).

(G) Pearson correlation coefficient (PCC) of LCI-EGFP with GM130 in cells transfected with the indicated mutant.

(H) Representative fluorescence pH and -log([Ca^2+^]) maps in *CalipHluor2.0*-labeled lysosomes in coelomocytes in *lci-1*^+/-^ worms extrachromosomally expressing the indicated variant of LCI.

(I) Maximum average lysosomal Ca^2+^ concentration in the indicated genetic backgrounds. Lysosomes with <0.1µM [Ca^2+^] are considered to have 0.1µM [Ca^2+^].

(J) Distribution of lysosomes with the indicated Ca^2+^ concentration measured using *CalipHluor2.0* in the indicated genetic backgrounds.

Scale bar 5µm. Boxes and bars represent s.e.m. and standard deviation, respectively. ns, not significant (p>0.05); *p<0.05; **p<0.01; ***p<0.001 (one-way ANOVA with Tukey post hoc test).

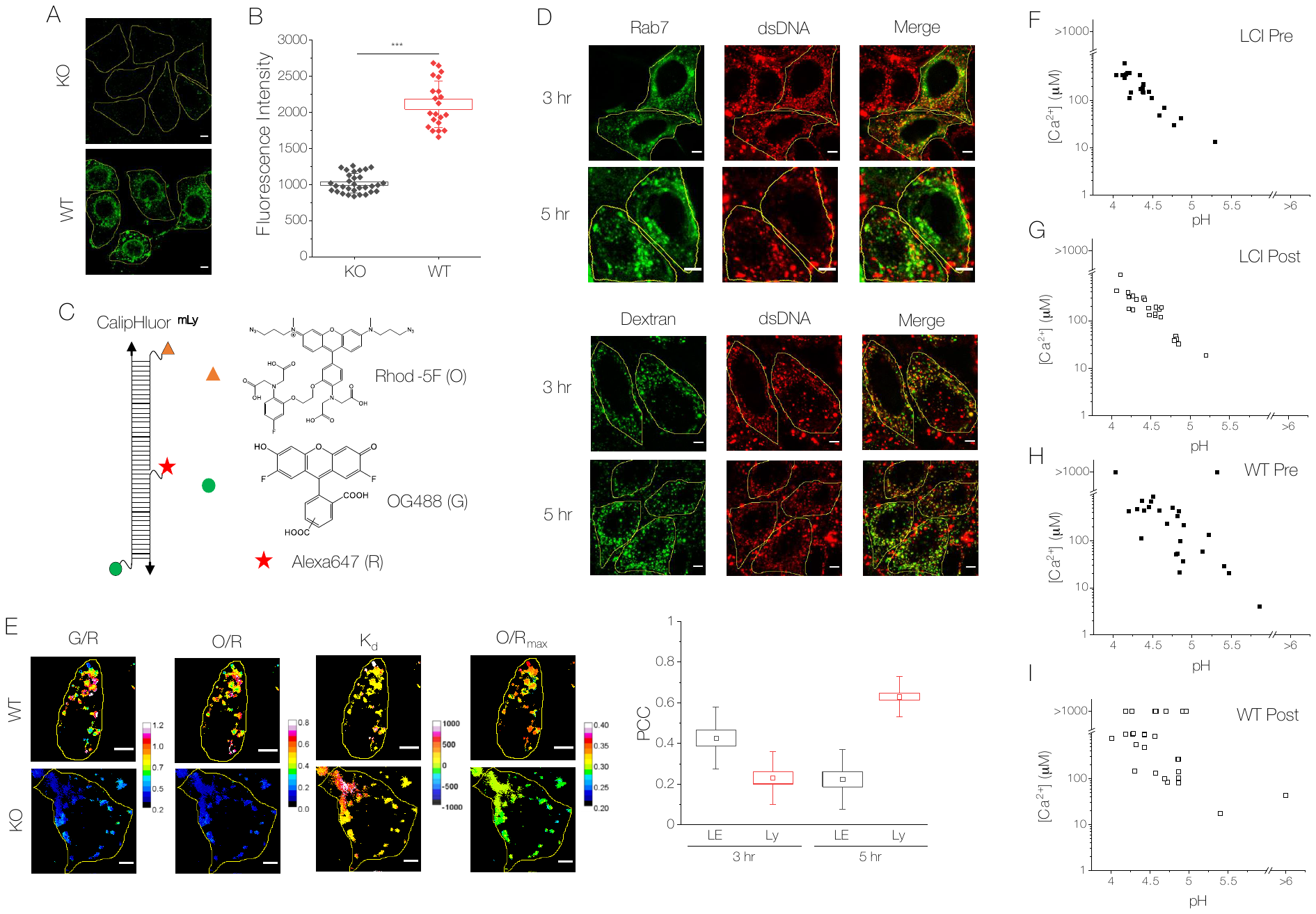

**Figure S5 | Justification and technology underlying single-lysosome Ca^2+^ measurements**.

(A) Representative immunofluorescence images showing endogenous expression of LCI in LCI KO HeLa cells and WT HeLa cells.

(B) Background-subtracted whole-cell fluorescence intensity of Alexa488-conjugated secondary antibody.

(C) Schematic of *CalipHluor^mLy^*, which consists of Rhod-5F (orange triangle), Oregon Green 488 (green circle), and Alexa647 (red star) on a DNA duplex.

(D) Top and middle, representative fluorescence images of HeLa cells pulsed with Alexa647-labeled dsDNA for 15 minutes and chased for the indicated amount of time. To look at late endosomal localization, cells were transfected with Rab7-GFP prior to dsDNA labeling (top). To look at lysosomal localization, cells were pulsed with FITC-dextran for 1 hour and chased overnight prior to dsDNA labeling (middle). Bottom, Pearson correlation coefficient (PCC) of dsDNA with endolysosomal markers at the indicated time points.

(E) Representative pseudocolored maps of the DCF/Alexa647 ratio (G/R) and Rhod-5F ratio (O/R) of *CalipHluor2.0* in WT and LCI KO HeLa cells. These maps are used according to equations in the Methods section to generate K_d_ and O/R_max_ maps.

(F-I) Individual 2-IM maps of lysosomal Ca^2+^ and pH in LCI KO HeLa cells before (F) and after (G) ATP addition, and in WT HeLa cells before (H) and after (I) ATP addition. Lysosomes below above O/R_max_ or G/R_max_ of *CalipHluor^mLy^* are shown as >1000µm Ca^2+^ or >pH 6, respectively.

Scale bar 5 µm. ***p<0.001

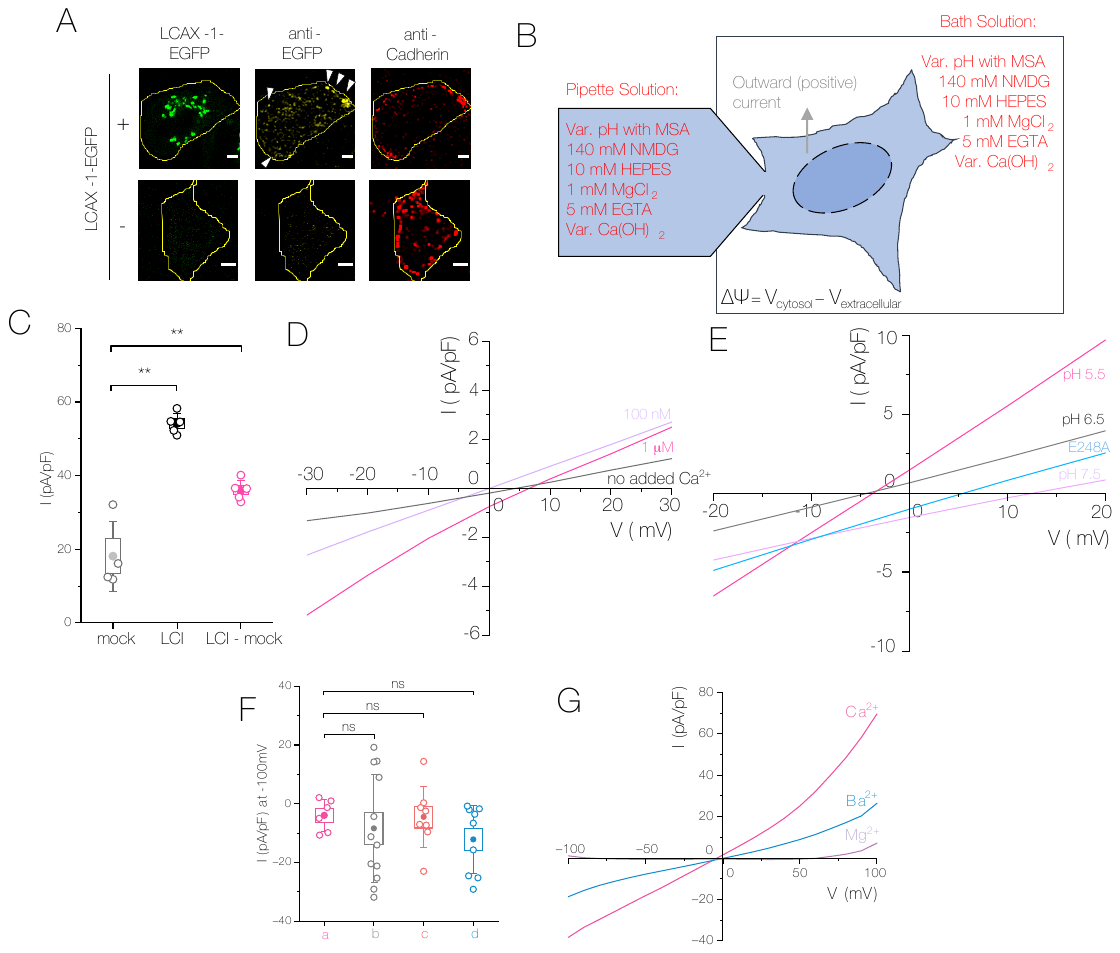

**Figure S6 | Electrophysiological characterization of LCI on the plasma membrane.**

(A) Representative image of LCI-EGFP-transfected HeLa cells and mock-transfected HeLa cells after fixation and immunostaining for EGFP (yellow) and cadherin (red), without permeabilization.

(B) Schematic representation of buffers used for whole-cell patch-clamping, and sign conventions used for current and membrane potential.

(C) Current density at +100mV across the plasma membrane of the indicated cell types under the indicated ionic conditions from Figure 7B.

(D) Zoomed-in version of Figure 7D to show reversal potentials under the indicated conditions.

(E) Zoomed-in version of Figure 7E to show reversal potentials under the indicated conditions.

(F) Current density at -100mV across the plasma membrane of the indicated cell types under the indicated ionic conditions from Figure 7G.\

(G) Average current density of mock-subtracted LCI current in HeLa cells under the indicated conditions, with the indicated divalent cation.

Boxes and bars represent s.e.m. and standard deviation, respectively. **p<0.01 (one-way ANOVA with Tukey post hoc test).

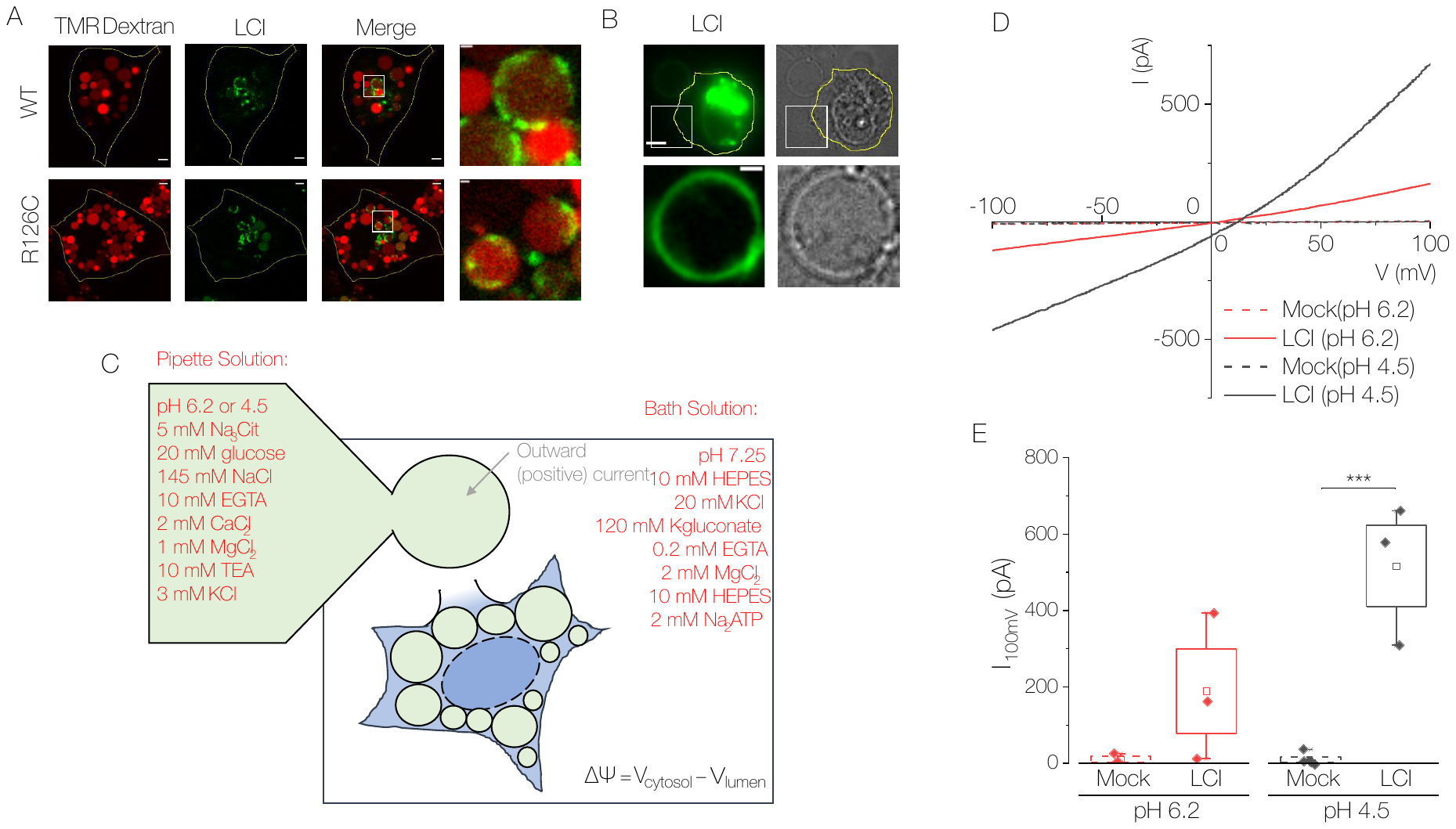

**Figure S7 | Electrophysiological characterization of LCI on isolated lysosomes**.

(A) Representative confocal images of COS-7 cells transiently transfected with WT or R126C LCI-EGFP (green). Lysosomes were labelled with 1 mg/mL TMR-dextran (red) and swollen with 5µM vacuolin-1 before imaging.

(B) Representative fluorescence of LCI-EGFP (green) on an isolated lysosome of COS-7 cells following overnight treatment with 5 µM vacuolin-1 and cell rupture.

(C) Schematic representation of buffers used for lysosome patch-clamping, and sign conventions used for current and membrane potential.

(D) Representative IV curves of lysosome patch-clamping using a ramp from -100mV to +100mV with buffers indicated in (C). Lysosomes of mock-transfected HeLa cells (dashed lines) or cells transfected with LCI-EGFP (solid lines) were patched with pipette solution of pH 6.2 (red) or pH 4.5 (black).

(E) Current at +100mV across the lysosome membrane from indicated cells with the indicated lysosomal pH.

Scale bar 5µm. Inset scale bar 2µm. Boxes and bars represent s.e.m. and outliers, respectively. *p<0.05.

**Supplementary Tables**

**Supplementary Table 2**

Internal symmetry of LCI.

| **Transmembrane Domains** | **Identity (%)** | **Similarity (%)** |
| --- | --- | --- |
| TMD1-TMD4 | 28.0 | 76.0 |
| TMD2-TMD5 | 35.0 | 65.0 |
| TMD3-TMD6 | 21.4 | 64.0 |

**Supplementary Table 3**

Internal symmetry of vcx-1.

| **Transmembrane Domains** | **Identity (%)** | **Similarity (%)** |
| --- | --- | --- |
| TMD1-TMD6 | 31.2 | 75.0 |
| TMD2-TMD7 | 35.0 | 70.0 |
| TMD3-TMD8 | 31.6 | 57.9 |
| TMD4-TMD9 | 26.7 | 66.7 |
| TMD5-TMD10 | 38.5 | 84.6 |

**Supplementary Table 4**

Templates used for homology-based modelling of LCI.

| **LCI Domain** | **Template Protein** | **Organism** | **Identity (%)** | **Similarity (%)** |
| --- | --- | --- | --- | --- |
| TMD-Reg | C-C chemokine receptor type 9 | *Homo sapiens* | 71.40 | 78.60 |
| TMD1 | VCX1 M2b | *Saccharomyces cerivisiae* | 36.36 | 54.54 |
| TMD2 | Monovalent cation-H^+^ antiporter | *Pyrococcus furiosus* | 85.70 | 85.70 |
| TMD3 | Two-pore calcium channel protein 2 | *Homo sapiens* | 50.00 | 70.00 |
| TMD4 | Calcium permeable stress-gated cation channel 1 | *Arabidopsis thaliana* | 61.50 | 76.90 |
| TMD5 | Calcium homeostasis modulator protein 2 | *Homo sapiens* | 44.40 | 55.60 |
| TMD6 | Neimann-Pick C1 protein | *Homo sapiens* | 63.20 | 78.90 |

**Supplementary Table 5**

Reversal potentials expected for exchanger with pipette buffer of pH 7.5 and 1µM Ca and bath buffer of 100 µM Ca and indicated pH.

| **Bath pH** | **Stoichiometry of Ca^2+^:H^+^** | **Reversal potential** |
| --- | --- | --- |
| 5.5 | 1:1 | -5.20E-14mV |
|  | 1:2 | ∞ |
|  | 1:3 | 234mV |
|  | 1:4 | 175.5mV |
|  | 1:5 | 156mV |
|  | 2:1 | 39mV |
|  | 2:3 | -117mV |
|  | 2:5 | 351mV |
|  | 3:1 | 46.8mV |
|  | 3:2 | 29.25mV |
|  | 3:4 | -58.5mV |
|  | 3:5 | -234mV |
| 6.5 | 1:1 | 58.5mV |
|  | 1:2 | ∞ |
|  | 1:3 | 58.5mV |
|  | 1:4 | 58.5mV |
|  | 1:5 | 58.5mV |
|  | 2:1 | 58.5mV |
|  | 2:3 | 58.5mV |
|  | 2:5 | 58.5mV |
|  | 3:1 | 58.5mV |
|  | 3:2 | 58.5mV |
|  | 3:4 | 58.5mV |
|  | 3:5 | 58.5mV |
| 7.5 | 1:1 | 117mV |
|  | 1:2 | ∞ |
|  | 1:3 | -117mV |
|  | 1:4 | -58.5mV |
|  | 1:5 | -39mV |
|  | 2:1 | 78mV |
|  | 2:3 | 234mV |
|  | 2:5 | -234mV |
|  | 3:1 | 70.2mV |
|  | 3:2 | 87.75mV |
|  | 3:4 | 175.5mV |
|  | 3:5 | 351mV |

**Supplementary Notes**

**Note S1**

**Localization of WT LCI.** Our data show that WT LCI is largely localized to the Golgi in HeLa cells, with minimal colocalization with TMR-dextran. However, previous experiments have found LCI in lysosome fractions[^1^](https://sciwheel.com/work/citation?ids=6612530&pre=&suf=&sa=0&dbf=0). In addition, we have detected overexpressed and endogenous LCI on membranes of lysosomes (Figure 2E and S7A-B)**.** Even a small fraction of LCI present on the lysosomes would be highly active, given the higher pH gradient there than across the Golgi membrane. Thus, the low lysosome localization does not preclude LCI from having a physiologically relevant role on the lysosome membrane.

**Note S2**

**Lysosomal calcium measurements.** The O/R ratio of ~50% of lysosomes of *lci-1*^+/-^ worms was below the O/R_min_ of *CalipHluor2.0*, indicating a Ca^2+^ concentration <100 nM that is not quantifiable by our probe (Figure 3E). Conversely, the O/R ratio of ~50% of lysosomes of *lci-1^+/-^* worms expressing WT human LCI was above the O/R_max_ of *CalipHluor2.0*, indicating a Ca^2+^ concentration >1 mM that is not quantifiable by our probe (Figure 3E). Thus, our reported effect of LCI on lysosomal Ca^2+^ levels in worms is actually an underestimation. This is why we only include quantifiable data points in statistics reported in Figure 3D, and do not report statistics in Figure S4I or Figure 5C.

Similarly, the O/R ratio of over 60% of lysosomes of LCI KO HeLa cells was below the O/R_min_ of *CalipHluor^mLy^* (Figure 5E**)***.* Only about 30% of lysosomes of WT HeLa cells had an O/R below the O/R_min_ (Figure 5E). Thus, the reported effect of LCI on lysosomal Ca^2+^ levels in cells is also an underestimation.

The lysosomal Ca^2+^ measurements of worms expressing mutants of human LCI are complicated by the heterozygous knockout background. Specifically, we see a surprisingly high level of lysosomal Ca^2+^ in *lci-1*^+/-^ worms expressing the G304R, E108A, and E248A mutants of human LCI. Yet, in all other assays, these mutants impair lysosomal Ca^2+^ import. Thus, it is likely that LCI acts as a dimer, and that the remaining copy of endogenous worm *lci-1* can dimerize with the mutant human LCI, and form a partially functional transporter. Given the smaller size of LCI compared to other Ca^2+^ transporters and exchangers, we hypothesize that it acts as a dimer.

**Note S3**

**Homologous regions of LCI.** The regions of LCI that show homology to vcx1 offer clues to how LCI may transport Ca^2+^ with high capacity, dependent on the lysosomal pH gradient. In vcx1, the proton motive force across the vacuole drives a conformational change where active site glutamate residues face the cytosol and maintain a negative charge[^2^](https://sciwheel.com/work/citation?ids=841504&pre=&suf=&sa=0&dbf=0). Under conditions of high cytosolic Ca^2+^, as seen in our experiments in Figure 4C in yeast and Figures 5A and 6 in mammalian cells, Ca^2+^ ions are coordinated by the cytosolic acidic helix to bring them near the active site. Coordination by the active site displaces water molecules to move helix M2b (designated in yellow in Figure 4A-B) towards the active site. This movement closes the cytosolic vestibule lined by M7b (designed in purple in Figure 4A-B) and opens a vacuolar cleft, such that the acidic pH of the vacuole lowers the Ca^2+^ affinity of active site glutamate residues and leads to release of Ca^2+^. This cyclical pumping occurs because of flexible helices around the active site and more rigid piston-like helices further away from the pore. Given that LCI possesses two Ca^2+^-binding sequences near regions homologous to the flexible internal helices of vcx1 and an acidic cytosolic helix in proximity, it follows that LCI functions similarly in response to high cytosolic Ca^2+^. However, the fact that LCI is much smaller than vcx1 and most other exchangers may indicate that it functions differently, for example as a dimer or as a pH-activated transporter instead of an exchanger.

**Note S4**

**Yeast color change.** Strains of *S. cereivisiae* that have mutations in certain steps of the adenine biosynthetic pathway (such as the ade2-1 mutation in K665[^3^](https://sciwheel.com/work/citation?ids=2569232&pre=&suf=&sa=0&dbf=0)) accumulate an adenine-intermediate-derived red pigment inside vacuoles[^4^](https://sciwheel.com/work/citation?ids=7548794&pre=&suf=&sa=0&dbf=0). The intermediate phosphoribosylaminoimidazole (AIR) is transported glutathione-dependently into vacuoles where it is polymerized and modified to form the characteristic red pigment. The structure of this pigment has yet to be fully established. Importantly, development of red pigmentation in ade2 mutants requires normal vacuolar function[^5^](https://sciwheel.com/work/citation?ids=7548816&pre=&suf=&sa=0&dbf=0). This concept has even been used to screen for chemicals that disrupt vacuolar function by loss of red pigmentation[^6^](https://sciwheel.com/work/citation?ids=7552505&pre=&suf=&sa=0&dbf=0). Interestingly, we see that K665 colonies grown on SD-Leu plates do not exhibit red pigmentation, but that LCI-transformed K665 colonies appear red. This red pigmentation is lost when plated on plates with high Ca^2+^. This implies that human LCI rescues vacuolar dysfunction in K665 under normal osmotic conditions, but that high Ca^2+^ causes vacuolar dysfunction even as LCI rescues lethality. While we cannot rule out the effect of LCI on other aspects of the adenine-derived pigment biosynthetic pathway, its lysosomal roles in humans and nematodes established elsewhere in this manuscript support the hypothesis that it rescues vacuolar dysfunction here.
